# The Sensitivity of ^13^C-Metabolic Flux Analysis to Biomass Composition Underscores the Need for Accurate Biomass Formulation in *Escherichia coli*

**DOI:** 10.64898/2026.10.21.760808

**Authors:** Yijing Cai, Maciek R. Antoniewicz

**Affiliations:** Department of Chemical Engineering, University of Michigan, Ann Arbor, Michigan 48109, USA

## Abstract

Quantifying intracellular fluxes through ^13^C-metabolic flux analysis (^13^C-MFA) provides a detailed view of cellular metabolism. While model formulation, including biomass synthesis reactions, is known to influence flux estimations, the degree to which biomass composition affects ^13^C-MFA outcomes in *Escherichia coli* remains debated. Here, we reconstructed a genome-scale ^13^C-MFA model of *E. coli* incorporating explicit nucleotide biosynthesis and a modular biomass formulation that separates protein, RNA, and other components. Using isotopic labeling experiments and systematic perturbations of biomass composition, we demonstrate that ^13^C-MFA can robustly differentiate subtle shifts in the protein/RNA ratio, with deviations as small as 2–3% producing statistically significant but biologically interpretable changes in model fit. Reaction-level analyses revealed consistent flux adjustments across amino acid and nucleotide biosynthesis, while pathways in one-carbon metabolism, particularly serine, glycine, and threonine interconversions, exhibited the strongest responses. Collectively, these results highlight the robustness and discriminative power of ^13^C-MFA and also demonstrate the importance of accurate biomass composition measurement in *E. coli* ^13^C-MFA, given the method’s intrinsic sensitivity to macromolecular stoichiometry.

## 1. Introduction

Quantifying intracellular flux distributions provides critical insights into cellular metabolism and regulatory mechanisms. Among available techniques, ^13^C-metabolic flux analysis (^13^C-MFA) stands out for its precision and ability to resolve fluxes that are otherwise inaccessible through transcriptomics, proteomics, or metabolomic^1–3^. This method combines isotopic tracer experiments with computational modeling: cells are cultured on ^13^C-labeled substrates, isotopic labeling patterns in metabolites (e.g., amino acids, organic acids) are quantified using MS or NMR, and fluxes are estimated by fitting the measured isotopomer distributions to a metabolic network model via iterative least-squares regression, as implemented in Metran^4^. The model’s goodness-of-fit is evaluated by the sum of squared residuals (SSR), and flux confidence intervals are derived from sensitivity analyses of the objective function^5^.

Since ^13^C-MFA relies on a defined metabolic model, its accuracy depends critically on both the network structure and the formulation of the biomass synthesis reaction, which represents the stoichiometric demands of macromolecular components (proteins, RNA, lipids, DNA, etc.). In most existing *E. coli* models, nucleotide biosynthesis pathways are omitted, and their contributions are instead lumped into the overall biomass reaction using precursors such as ribose-5-phosphate or phosphoenolpyruvate^6,7^. Although this simplification allows model convergence, it can obscure the influence of nucleotide metabolism and biomass composition on flux estimations.

While the importance of accurate biomass composition has long been recognized in flux balance analysis (FBA)^8,9^, it is often overlooked in ^13^C-MFA. A survey of the Web of Science database identified over 700 studies employing ^13^C-MFA, yet only 16 explicitly measured or discussed biomass composition. There have been rising concerns regarding the importance of elucidating biomass composition when dealing with different organisms and environments, such as yeast^10^, *Rhodococcus opacus*^11^, photosynthetic microorganisms^12^, etc. in recent years. But there are counter arguments regarding its importance when studying with *E. coli*. Gopalakrishnan et, al. reported and claimed that for *E. coli*, whose biomass composition are relatively consistent, ^13^C-MFA results are largely insensitive to moderate perturbations in each biomass component and show no significant flux range shift, and the error is major coming from isotopic measurement^6^. Long et al. reported that *E. coli*’s protein and RNA composition can shift as high as 10% with single gene knockout^13^, and emphasized the importance of specifying biomass composition^2,14^.

In this study, we revisited this question by reconstructing the *E. coli* ^13^C-MFA model based on the latest genome-scale framework and explicitly expanding the biomass reaction. The lumped biomass flux was separated into three adjustable components: protein, RNA, and others (lipids, DNA, etc.), enabling direct simulation of altered biomass compositions based on experiment measurements^13^. Using this optimized model, we systematically evaluated ^13^C-MFA sensitivity under different protein/RNA ratios and overall biomass fractions. Our analysis reveals that ^13^C-MFA fitness and predicted flux distributions are highly sensitive to even small changes in protein and RNA composition, larger than 2-3% fluctuation will lead to unacceptable simulation results. Which proves the preciseness of isotopic labeling experiment and the sensitivity of ^13^C-MFA towards biomass composition. Notably, reactions involved in one-carbon metabolism, including serine, glycine and threonine interconversions, exhibit pronounced shifts, emphasizing the need for accurate biomass composition measurements in reliable ^13^C-MFA applications.

## 2. Results

### 2.1. Protein and RNA composition impact ^13^C-MFA fitness

To investigate the sensitivity of ^13^C-metabolic flux analysis (^13^C-MFA) to variations in biomass composition, we reconstructed the metabolic model of *E. coli* MG1655 based on the latest genome-scale model iML1515^15^. Specifically, we decomposed the lumped biomass synthesis reaction into distinct components: protein synthesis, nucleic acid synthesis, and other biomass-associated reactions. In previous models, nucleotide biosynthesis pathways were often oversimplified and directly incorporated into the biomass reaction. By explicitly modeling these reactions, our framework enables flexible adjustment of biomass composition inputs. The updated model was validated and produced results consistent with the original formulation (SI table 1).

We then applied ^13^C-MFA across a range of biomass composition scenarios—modulating the “other” biomass fraction (g_other) to 15%, 20%, 25%, and 30%, and varying protein and RNA mass fractions in 1% increments—while identifying the best-fit solution (lowest sum of squared residuals, SSR) for each condition (SI table 2). As shown in Figure 1A, the best model fits consistently occurred at a protein-to-RNA ratio of approximately 3, regardless of g_other. Deviations from this ratio, even by 2–3%, led to significant increases in SSR, often exceeding the acceptable confidence threshold. Only 5% difference of protein percentage set up is acceptable for simulation with g_other at 15% and 20%, and only 4% are acceptable when g_other at 25% and 30% (Figure 1B-D). These findings suggest that while ^13^C-MFA is relatively robust to changes in other biomass components, it is highly sensitive to the precise distribution between protein and RNA. The acceptable fluctuation range is much narrower than different *E. coli* strain’s potential biomass composition fluctuation^13^. This challenges prior claims that ^13^C-MFA results are largely insensitive to biomass composition fluctuations^6^.

**Figure 1:**
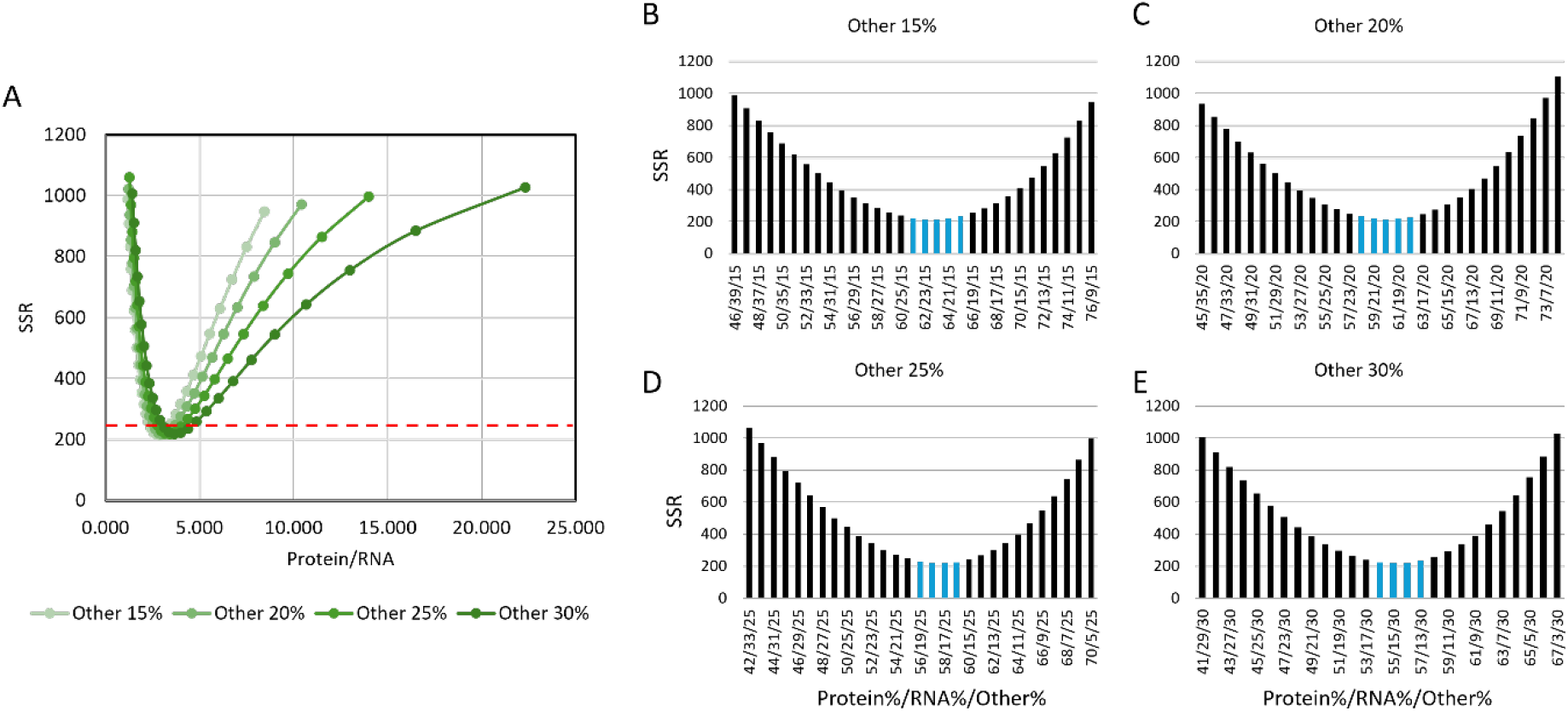
SSR landscape of ^13^C-MFA across biomass composition space. (A) SSR values for each Protein/RNA at fixed g_other (15%, 20%, 25%, 30%) from ^13^C-MFA simulations. The red dashed line marks the acceptable SSR threshold. Only simulations with SSR below the red dashed line indicate it’s accepted by ^13^C-MFA (B-E) Separated SSR values for each g_other setups. The blue bars were the simulations that is acceptable based on statistical verification of ^13^C-MFA

### 2.2. Reaction-Level Sensitivity to Biomass Composition Changes

To further evaluate the sensitivity of ^13^C-MFA simulations to biomass composition, we analyzed flux variability across individual reactions under different biomass settings (see Supplementary table 1). For each g_other scenario, we compared model-predicted fluxes from the best-fit protein/RNA composition to those obtained after a ±5% shift in protein/RNA content (Figure 2). The results show that over 50% of reactions experienced flux changes greater than 5%, and more than 30% of reactions varied by over 10%. Notably, ∼5% of reactions deviated by more than 50%, and ∼3% showed changes exceeding 100%.

**Figure 2:**
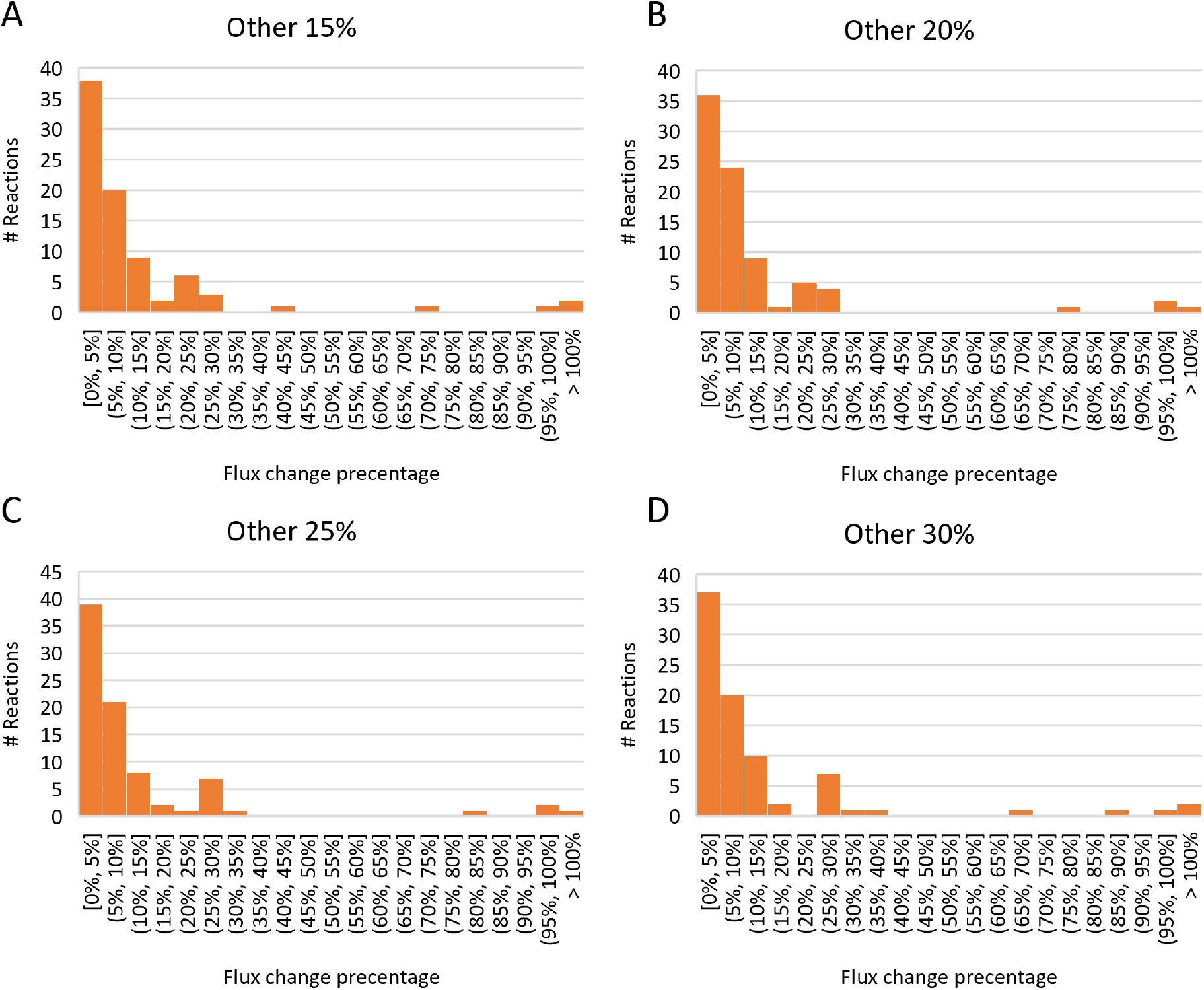
Distribution of flux-changes in response to biomass composition perturbation. Histograms show the distribution of flux deviation for all reactions under biomass shift (± 5% change in Protein or RNA composition) relative to the best-fit conditions at g_other of 15% (A), 20% (B), 25% (C), and 30% (D).

Despite large changes in g_other, its influence on most reactions remained minimal (Supplementary table 1). In contrast, altering the protein/RNA ratio had pronounced effects. Several reactions in glycolysis exhibited a positive correlation with increased protein composition, while reactions in the pentose phosphate pathway (PPP) displayed mixed responses with protein content. The TCA cycle, pyruvate metabolism, and acetate synthesis reactions were relatively unaffected by biomass composition changes. Most nucleotide biosynthesis reactions exhibited a negative correlation with increasing protein content, while the majority of amino acid biosynthetic reactions displayed a positive correlation—an expected result given the simultaneous decrease in RNA and increase in protein fractions (Supplementary material S1).

Interestingly, serine and glycine biosynthesis, along with glycine cleavage reactions, diverged from this trend and showed negative correlations with increasing protein composition (Figure 3A-C). This can be explained by the role of serine-derived glycine in generating 5,10-methylenetetrahydrofolate, a key one carbon donor critical for nucleotide, methionine, and histidine biosynthesis. When RNA demand is reduced, one carbon metabolism is predicted to be downregulated accordingly, leading to reduced flux through serine and glycine pathways. In contrast, glycine production from threonine exhibited a switch-like activation as protein content increased (Figure 3D). While this reaction is typically inactive in wild-type *E. coli*, its activation in the altered biomass scenarios suggests a compensatory mechanism in the model to meet the increased glycine demand from protein synthesis.

**Figure 3:**
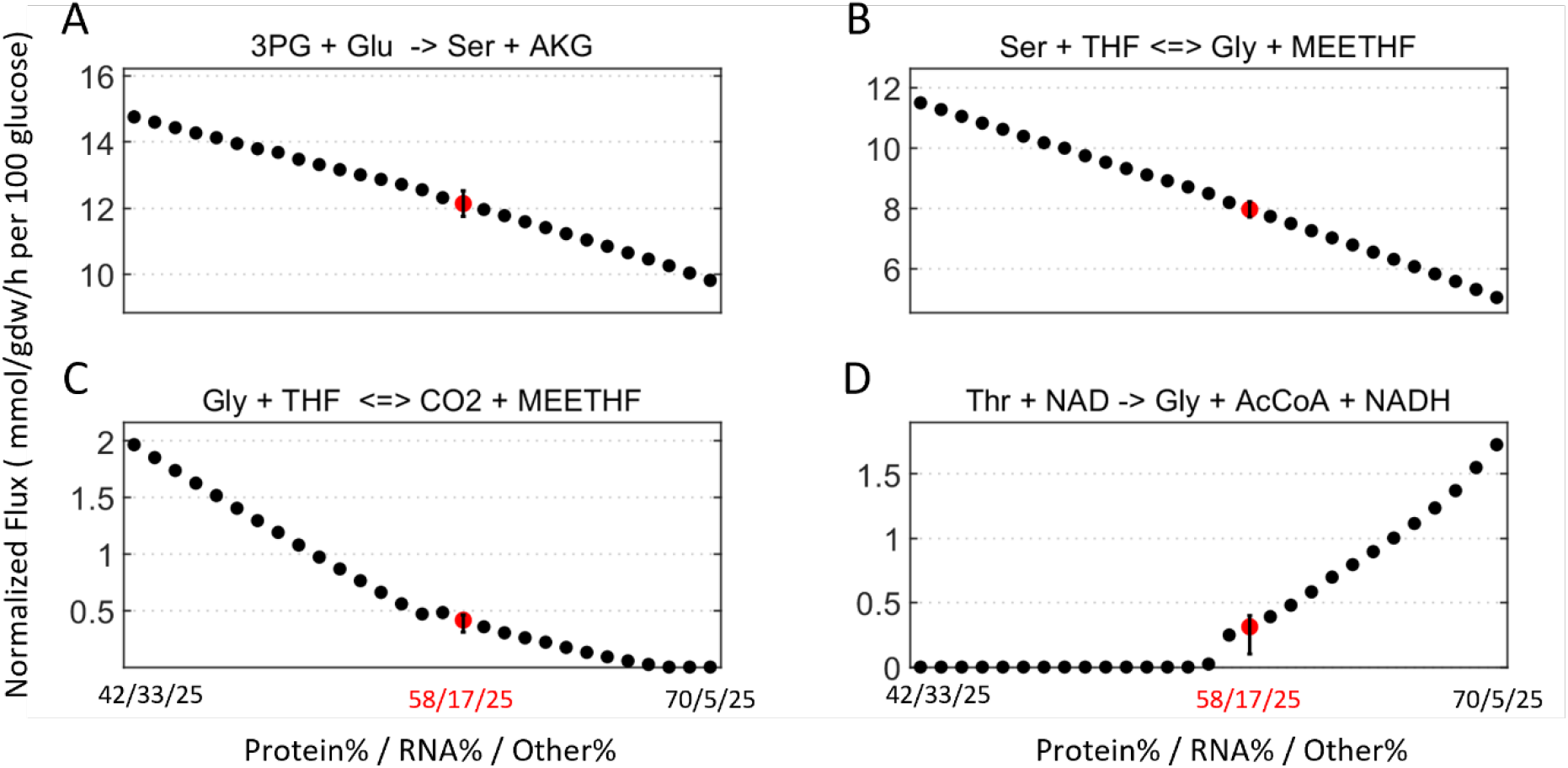
Sensitivity of one carbon metabolism reactions to biomass composition shifts. Flux predictions from ^13^C-MFA simulations under varying protein and RNA compositions (with g_other fixed at 25%) are shown for four key reactions. (A) 3PG + Glu → Ser + AKG (serine biosynthesis), (B) Ser + THF ⇌ Gly + METHF (serine hydroxymethyltransferase), (C) Gly + THF ⇌ CO_2_ + METHF (glycine cleavage system), (D) Thr + NAD → Gly + AcCoA + NADH (threonine-to-glycine pathway). Each black dot represents the flux under a specific biomass composition. The x-axis labels indicate the tested biomass compositions (protein/RNA/g_other). Red points with error bar indicate flux and standard deviation for the best-fit composition (58/17/25) determined by ^13^C-MFA.

These findings highlight the complex, non-linear relationships between biomass composition and intracellular flux predictions, particularly for one-carbon metabolism. Reactions involving serine, glycine, and nucleotide biosynthesis are especially sensitive to biomass composition assumptions. Without accurate, experimentally determined biomass parameters, ^13^C-MFA simulations risk producing misleading flux estimates in these pathways.

## 3. Discussion and conclusions

Our results demonstrate that 13C-MFA is highly sensitive to biomass composition, especially the ratio of protein to RNA. When this ratio deviated by as little as 2–3% from the best-fit condition, the model’s sum of squared residuals (SSR) increased sharply, often exceeding the acceptable threshold for a valid fit. This finding directly contradicts earlier claims that ^13^C-MFA is robust to biomass composition changes within a 10% range^6^. While the co-factors, DNA and other precursors from biomass composition (“g_other”) had relatively small influence, minor changes in protein and RNA fractions significantly affected flux estimations across central metabolism.

At the reaction level, flux predictions in glycolysis and the pentose phosphate pathway (PPP) were particularly sensitive to protein/RNA shifts, while the TCA cycle and acetate secretion fluxes remained more unaffected. Most amino acid biosynthesis and nucleotide pathways also showed consistent directional changes in response to composition shifts. In particular, serine and glycine biosynthesis, glycine cleavage, and threonine-to-glycine conversion were strongly impacted, reflecting the tight coupling of these reactions to one-carbon metabolism and RNA synthesis demands. Interestingly, the model predicted activation of the threonine-to-glycine pathway under high protein demand—a pathway typically inactive in wild-type E. coli—suggesting potential compensatory metabolic adaptation in response to increased glycine demand.

To evaluate the effect of biomass composition from a stoichiometric perspective, we also performed Flux Variability Analysis (FVA). In this setup, glucose uptake was fixed at 100 units, while acetate secretion was constrained between 45–75 mol per 100 mol glucose, and biomass yield was limited to 0.46 gDW/g ±5%. The g_other fraction was fixed at 25%, and protein/RNA composition varied across the same range as in the ^13^C-MFA simulations. FVA showed similar directional trends in flux changes across conditions, and the fluxes estimated by ^13^C-MFA generally fell within the predicted FVA ranges (Supplementary material S2).

However, FVA predicted narrow flux ranges for most amino acid biosynthesis and nucleotide pathways, while serine, glycine, and threonine-related reactions showed broad feasible ranges. This suggests that while FVA captures overall feasibility, it lacks the resolution to pinpoint specific pathway usage within the one-carbon network. In contrast, ^13^C-MFA, by leveraging isotopic labeling data, accurately resolved pathway preference, including the preference for glycine production from serine. The broader FVA range for glycolysis and PPP fluxes likely reflects partial uncertainty due to the unconstrained variability in acetate secretion within the allowed range.

Together, these findings emphasize that accurate biomass composition is critical for reliable ^13^C-MFA. One-carbon metabolism and nucleotide-related pathways are particularly vulnerable to small shifts in composition, and their accurate quantification requires isotopic labeling data. FVA alone, while helpful for bounding feasible flux distributions, cannot replace the resolution of 13C-MFA. As biomass composition can vary significantly in stressed, nutrient-limited, or genetically modified strains, we recommend that future MFA studies incorporate either measured biomass data or model structures that allow flexible adjustment of biomass demands, such as the modular biomass approach used here. This will help avoid misinterpretation of fluxes, especially in key regulatory and biosynthetic pathways.

## Methods and materials

### 4.1. Strains and culture conditions

*E. coli* K-12 BW25113 strain was used in this study. The strain was cultured aerobically in M9 minimal medium supplemented with 2 g/L glucose at 37°C overnight and then inoculated into 10 mL experiment medium with 1:100 ratios. The experiment cultures were cultivated in parallel with M9 with 2 g/L of glucose tracers, [1,6-^13^C]glucose and [1,2-^13^C]glucose.

Cell growth was monitored by measuring optical density at 600 nm (OD_600_) using an Eppendorf BioPhotometer. OD values were converted to cell dry weight concentrations using the standard factor of 1.0 OD_600_ = 0.3 gDW/L. Supernatant samples were collected every 2 hours for glucose and acetate quantification. Cell pellets were harvested at mid-exponential phase, washed with glucose-free M9 medium, and stored at −20 °C for downstream GC-MS analysis.

### 4.2. GC-MS Analysis

Gas chromatography–mass spectrometry (GC-MS) analysis was performed using an Agilent 7890A GC system equipped with an HP-5MS capillary column (30 m, 0.25 mm i.d., 0.25 μm film thickness; Agilent J&W Scientific) and connected to an Agilent 5977B Mass Spectrometer operating in electron impact (EI) mode at 70 eV. Helium was used as the carrier gas at 1 mL/min. Source, quadrupole, interface, and inlet temperatures were maintained at 230 °C, 150 °C, 280 °C, and 250 °C, respectively. Mass spectra were acquired in single ion monitoring (SIM) mode with 4 ms dwell time per ion.

Proteinogenic amino acid labeling was measured using TBDMS (tert-butyldimethylsilyl) derivatization as previously described^4^. Extracellular acetate was quantified via derivatization of a 1:1 mixture of culture supernatant and 4 mM [4-^2^H]acetate internal standard as described before ^16^. Glucose consumption was assessed using hydroxylamine hydrochloride and propionic anhydride derivatization of a 1:1 mixture of culture supernatant and [U-^13^C]glucose, following established protocols ^4^.

### 4.3. Metabolic Network Modeling and ^13^C-Metabolic Flux Analysis

^13^C-metabolic flux analysis (^13^C-MFA) was performed using the software Metran ^17^, which implements the elementary metabolite unit (EMU) framework ^18^. The base model includes major central carbon metabolism pathways (EMP, PPP, TCA cycle, glyoxylate shunt, and Entner–Doudoroff pathway), amino acid biosynthesis, a transhydrogenase reaction, methionine uptake, and a lumped biomass reaction. In this study, we extended the model to include nucleotide biosynthesis pathways and rewrote the biomass reaction to separate it into three adjustable compartments: protein, RNA, and other biomass components (g_other) (see Supplementary table 1).

Biomass composition was systematically varied by adjusting protein and RNA fractions in 1% increments, while g_other was tested at 15%, 20%, 25%, and 30%. For each condition, fluxes were estimated by minimizing the variance-weighted sum of squared residuals (SSR) between experimentally measured and simulated mass isotopomer distributions and acetate yields using nonlinear least-squares regression. All fluxes were normalized to the glucose uptake rate, which was determined from extracellular glucose consumption, acetate secretion, and growth rate measurements^4^.

The best-fit condition for each biomass setup was evaluated for statistical consistency using a χ^2^-test at the 95% confidence leve^19^. Confidence intervals for flux estimates were calculated based on SSR sensitivity analysis. All simulations and statistical analyses were conducted in MATLAB.

### 4.4. Flux Variability Analysis (FVA)

Flux variability analysis (FVA) was performed to evaluate the possible ranges of fluxes under different biomass composition scenarios. A MATLAB-based FVA script was used to calculate the upper and lower flux bounds while maintaining stoichiometric feasibility. The metabolic network used for FVA matched the model employed in ^13^C-MFA, including the addition of nucleotide biosynthesis pathways and a modular biomass formulation, where biomass was divided into protein, RNA, and other components (g_other).

In all FVA simulations, glucose uptake was fixed at 100 mmol/gDW/h. Acetate production was constrained between 45 and 75 mmol per 100 mmol glucose consumed, and biomass yield was fixed at 0.46 gDW/g with a ±5% tolerance. The g_other fraction was held constant at 25%, while protein and RNA compositions were varied systematically, consistent with the conditions tested in ^13^C-MFA.

FVA was performed using a linear programming solver in MATLAB. For each reaction, the minimum and maximum fluxes were calculated under the given constraints. The results were used to identify trends in flux flexibility and to compare with corresponding ^13^C-MFA flux estimates. The agreement or divergence between FVA-predicted ranges and ^13^C-MFA values was used to assess model sensitivity and the added resolution provided by isotopic labeling data.

## Supporting information

Supplementary Informations

## Notes

### Competing Interest Statement

The authors have declared no competing interest.

