## Supplementary Informations for "The Sensitivity of ^13^C-Metabolic Flux Analysis to Biomass Composition Underscores the Need for Accurate Biomass Formulation in *Escherichia coli*"

**Figure S1: Flux estimation of reactions from ^13^C-MFA across biomass compositions**

(A) **g_other 15%**

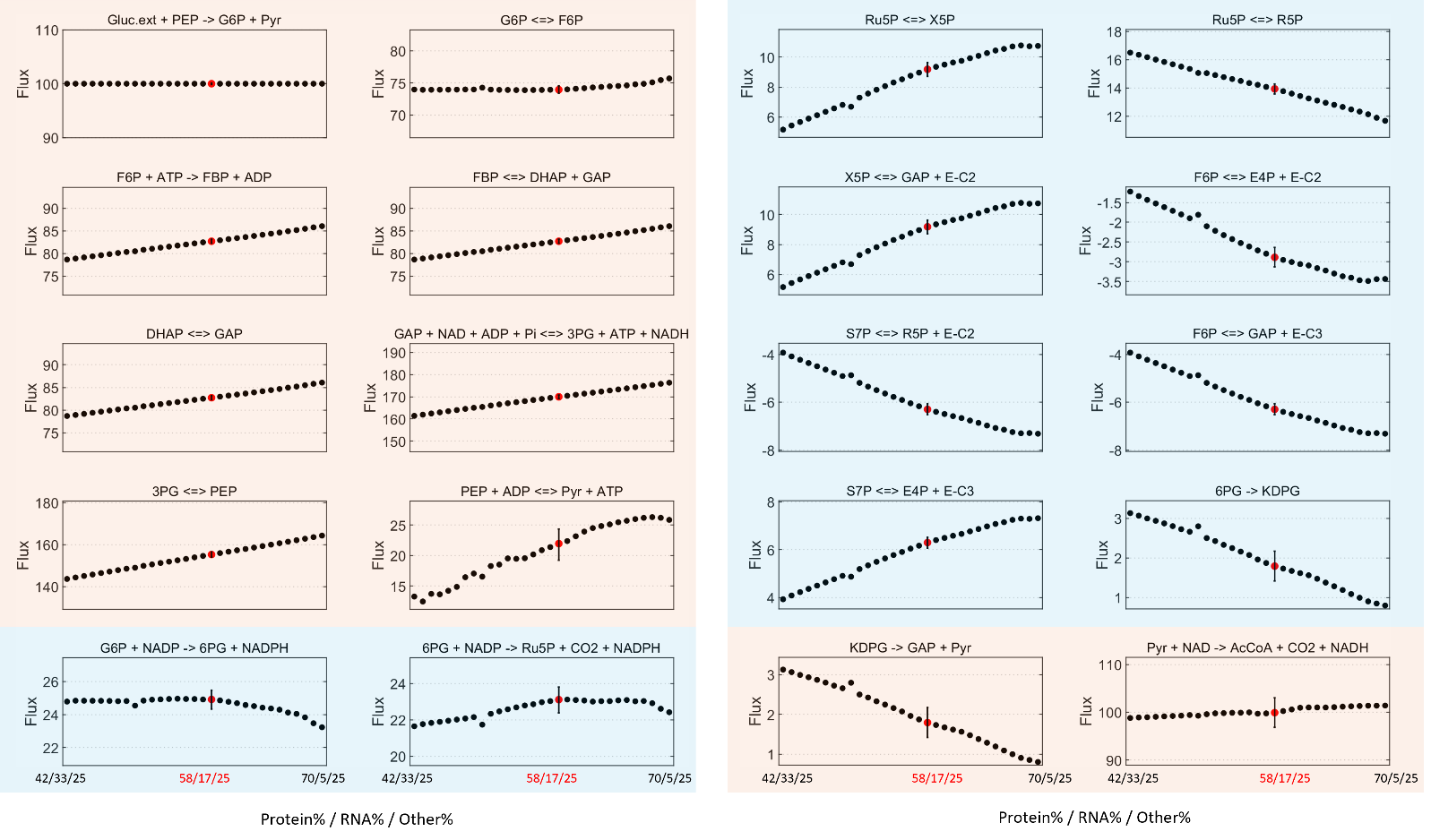

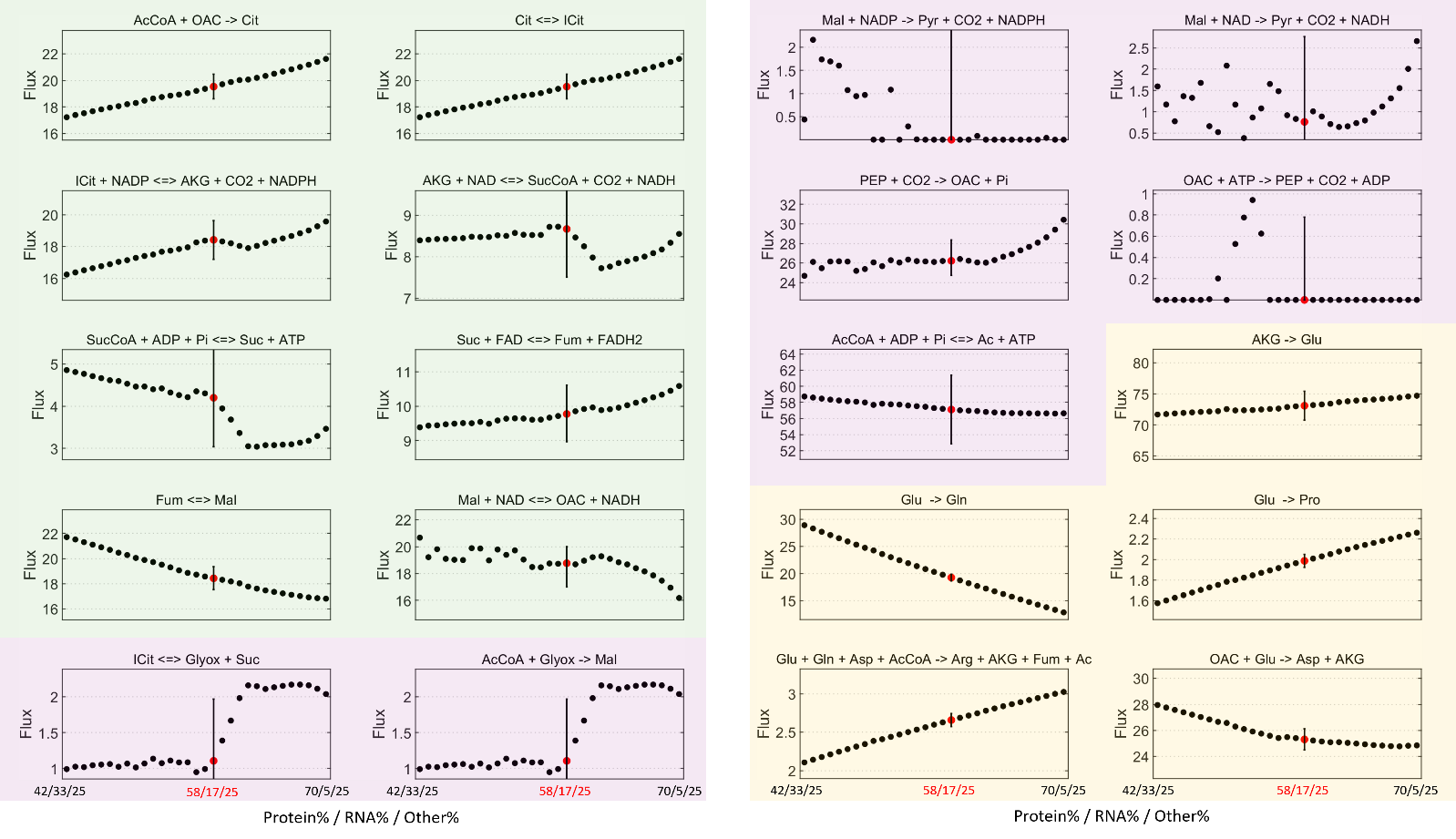

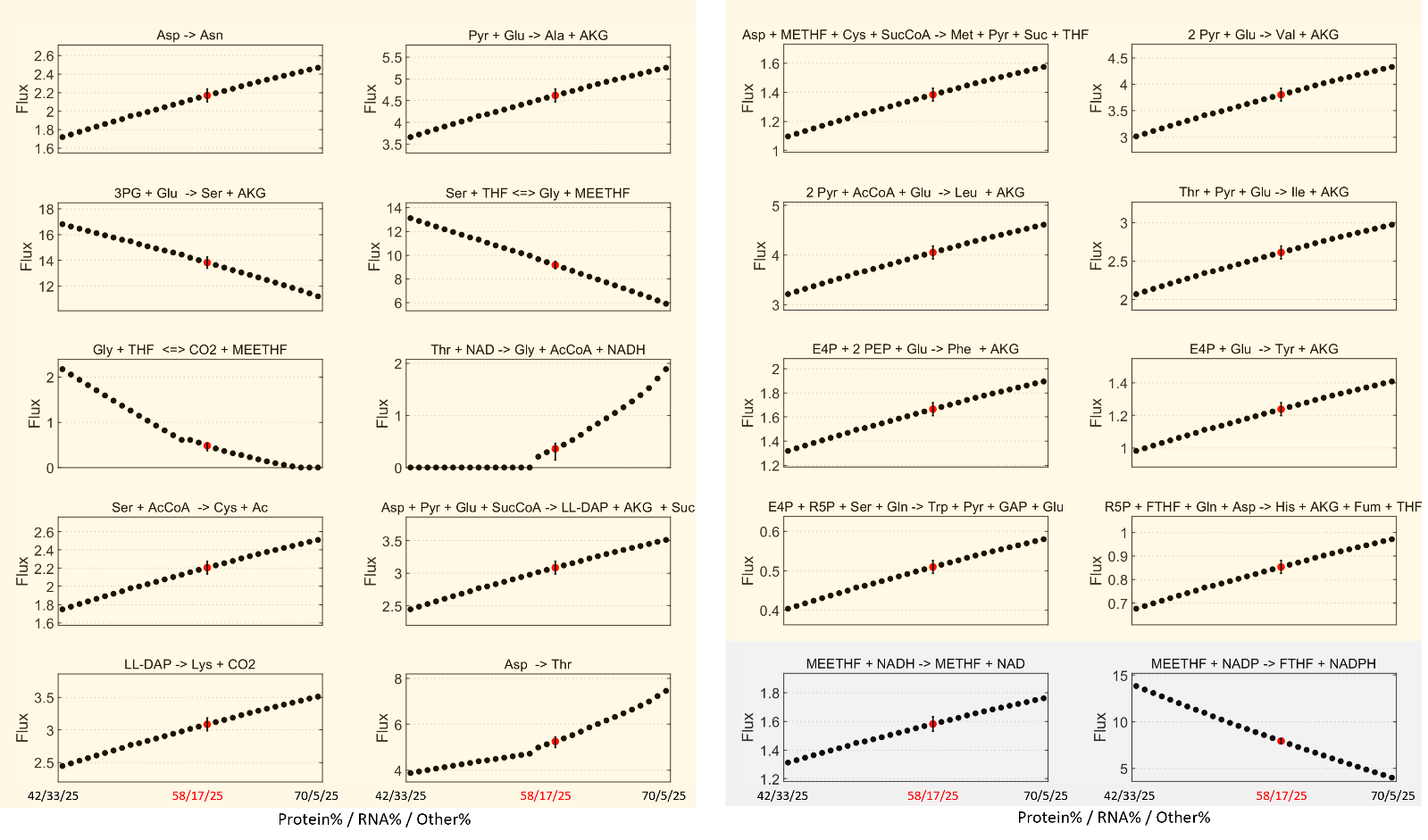

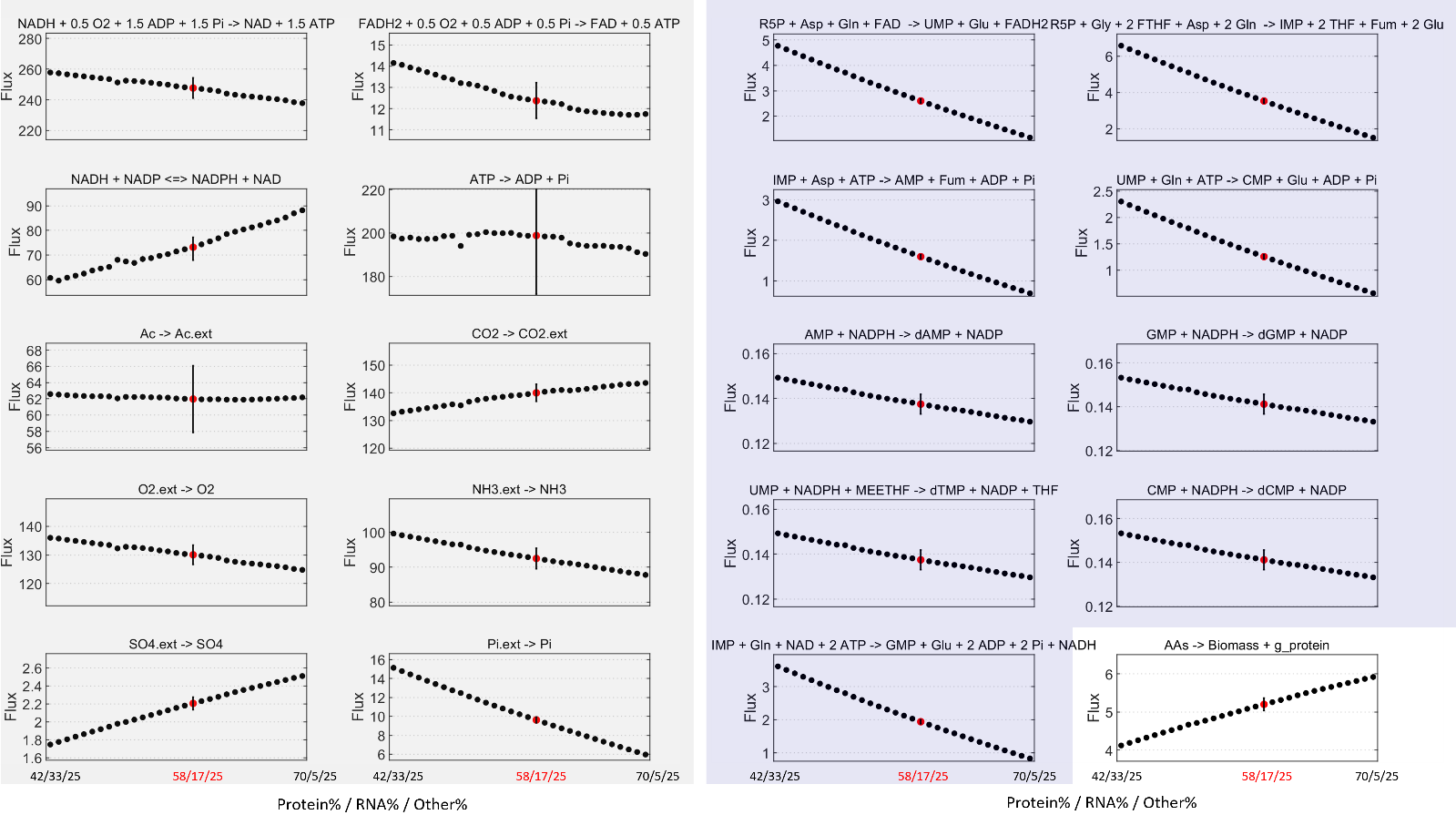

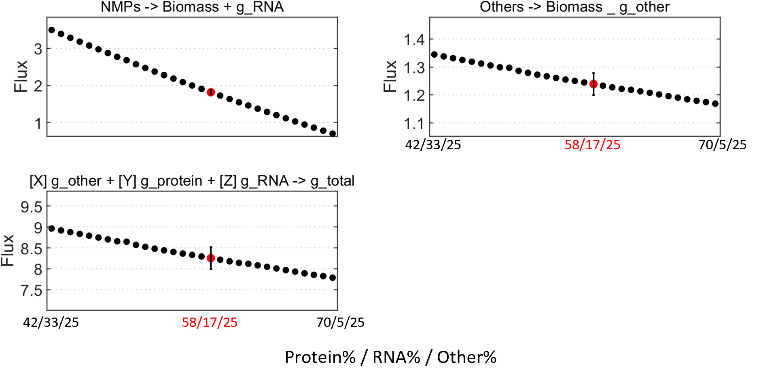

(B) **g_other 20%**

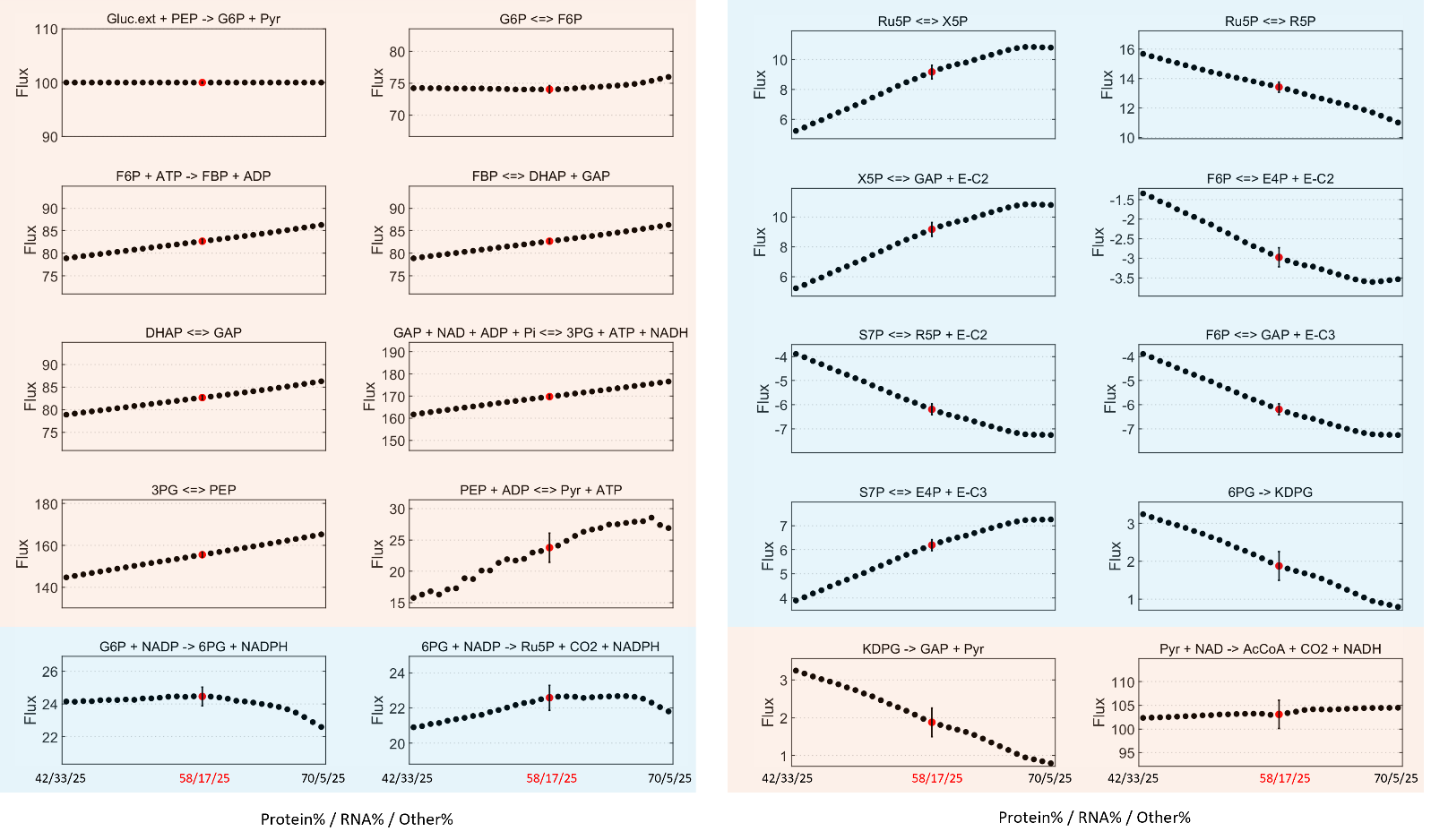

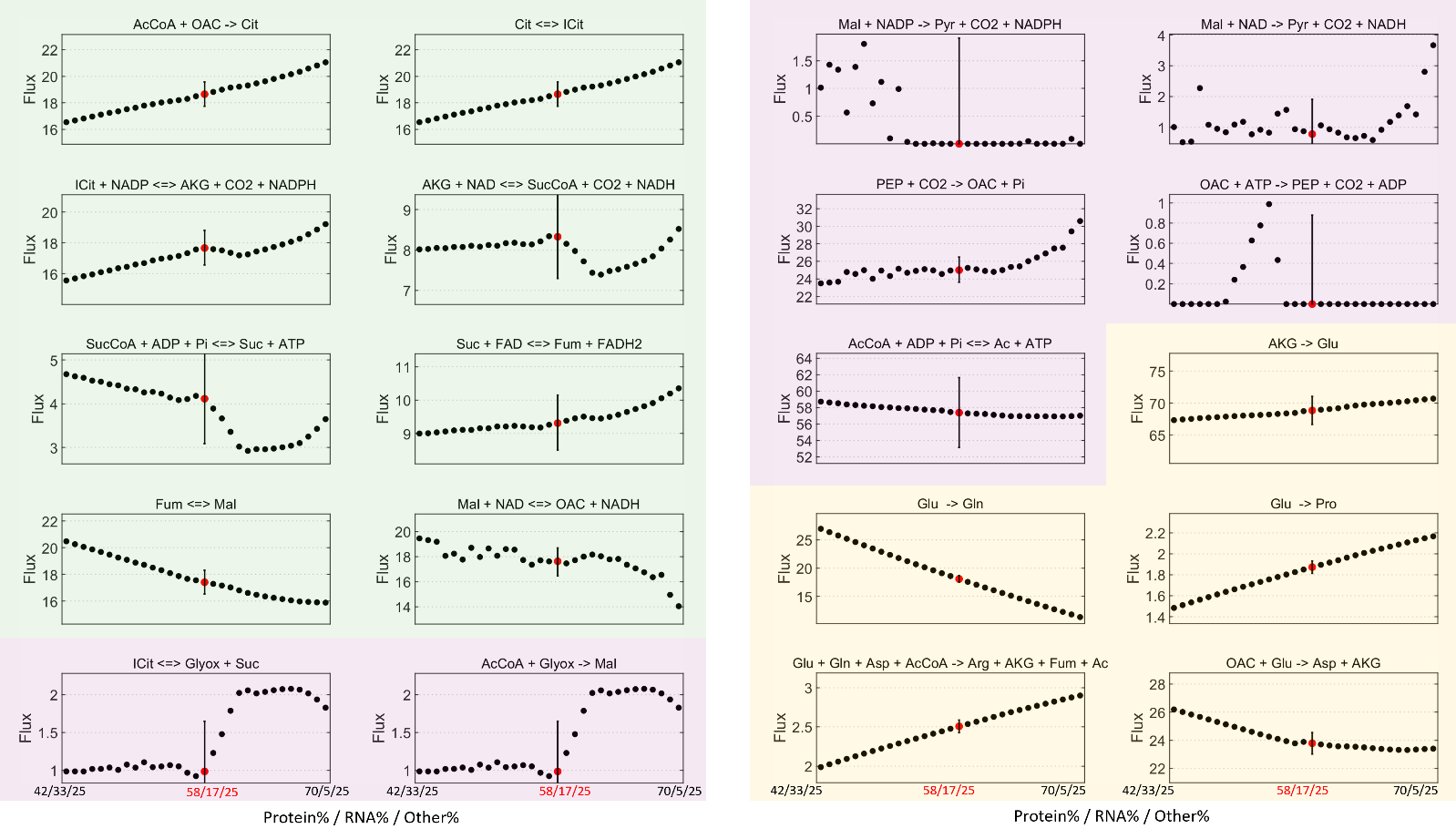

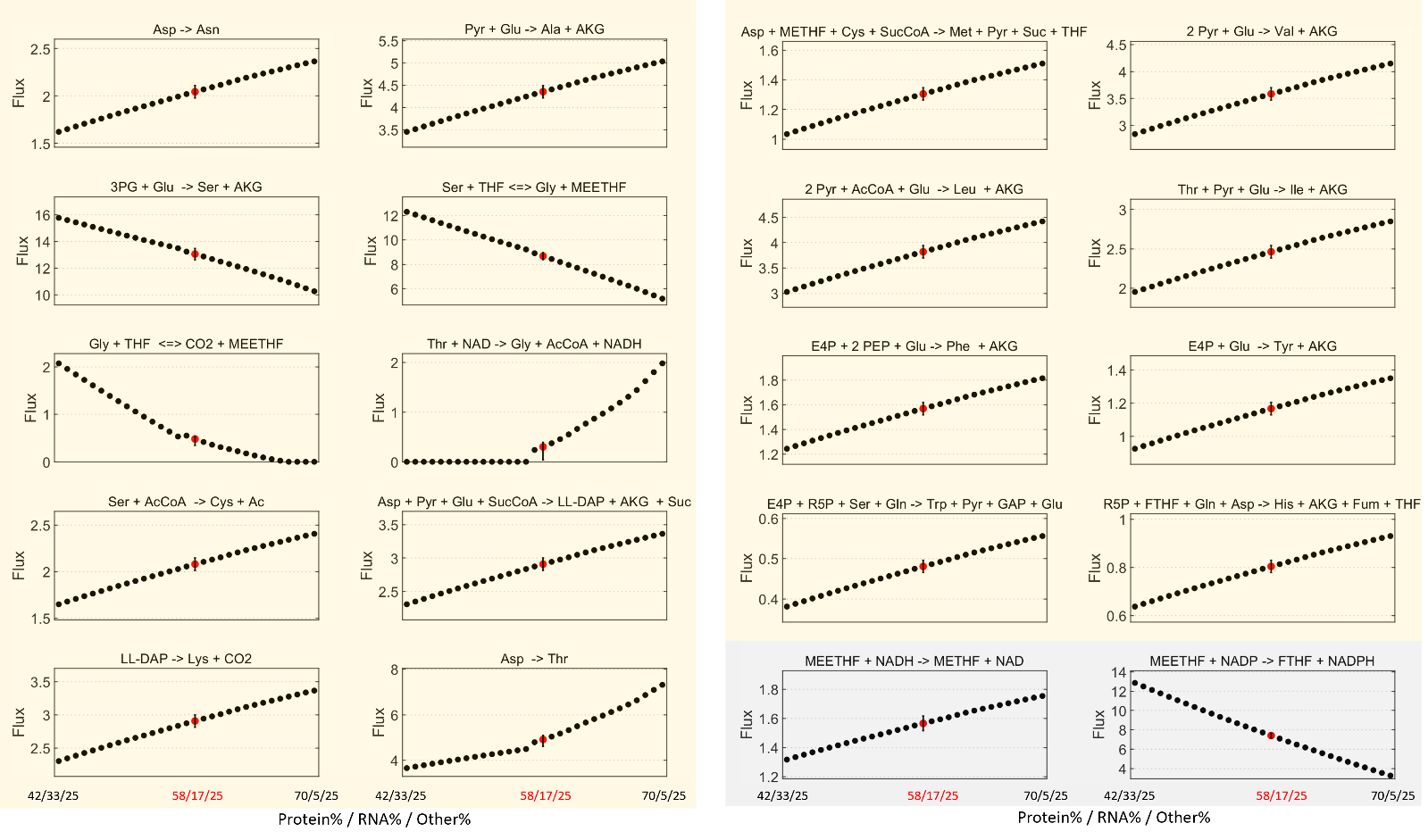

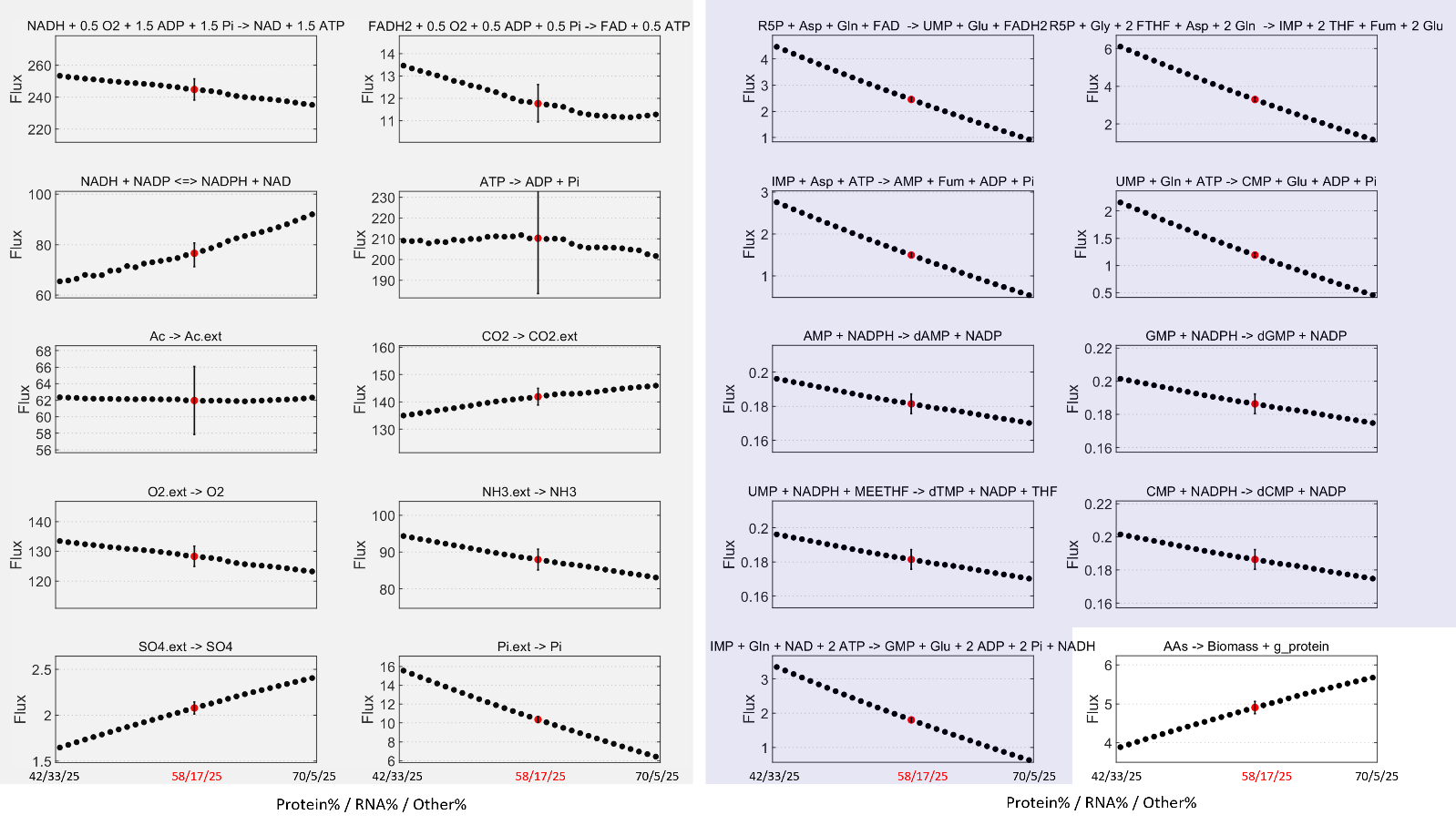

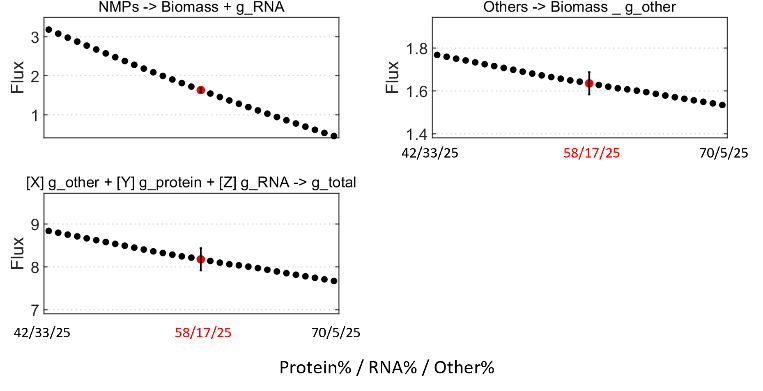

(C) **g_other 25%**

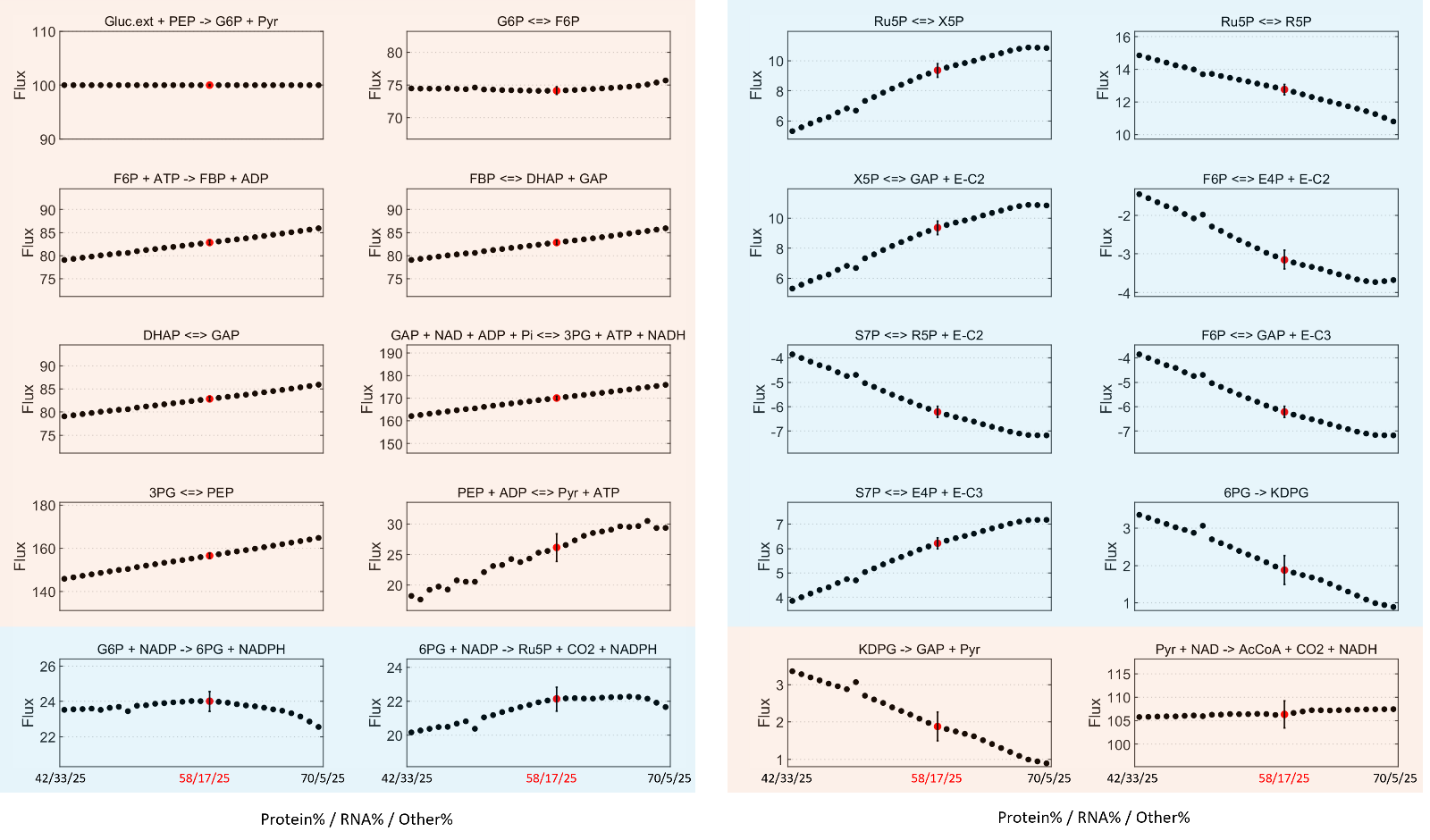

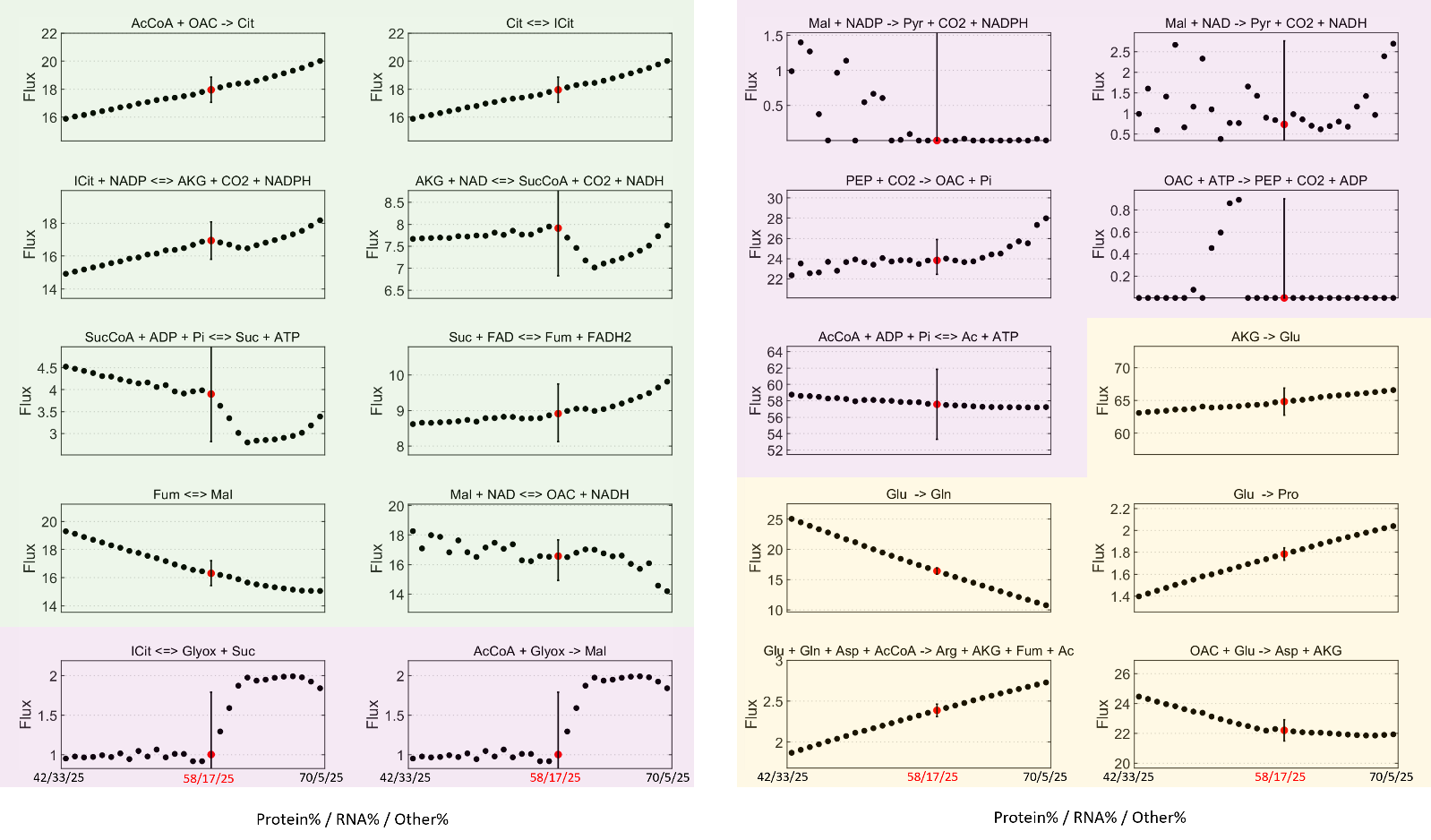

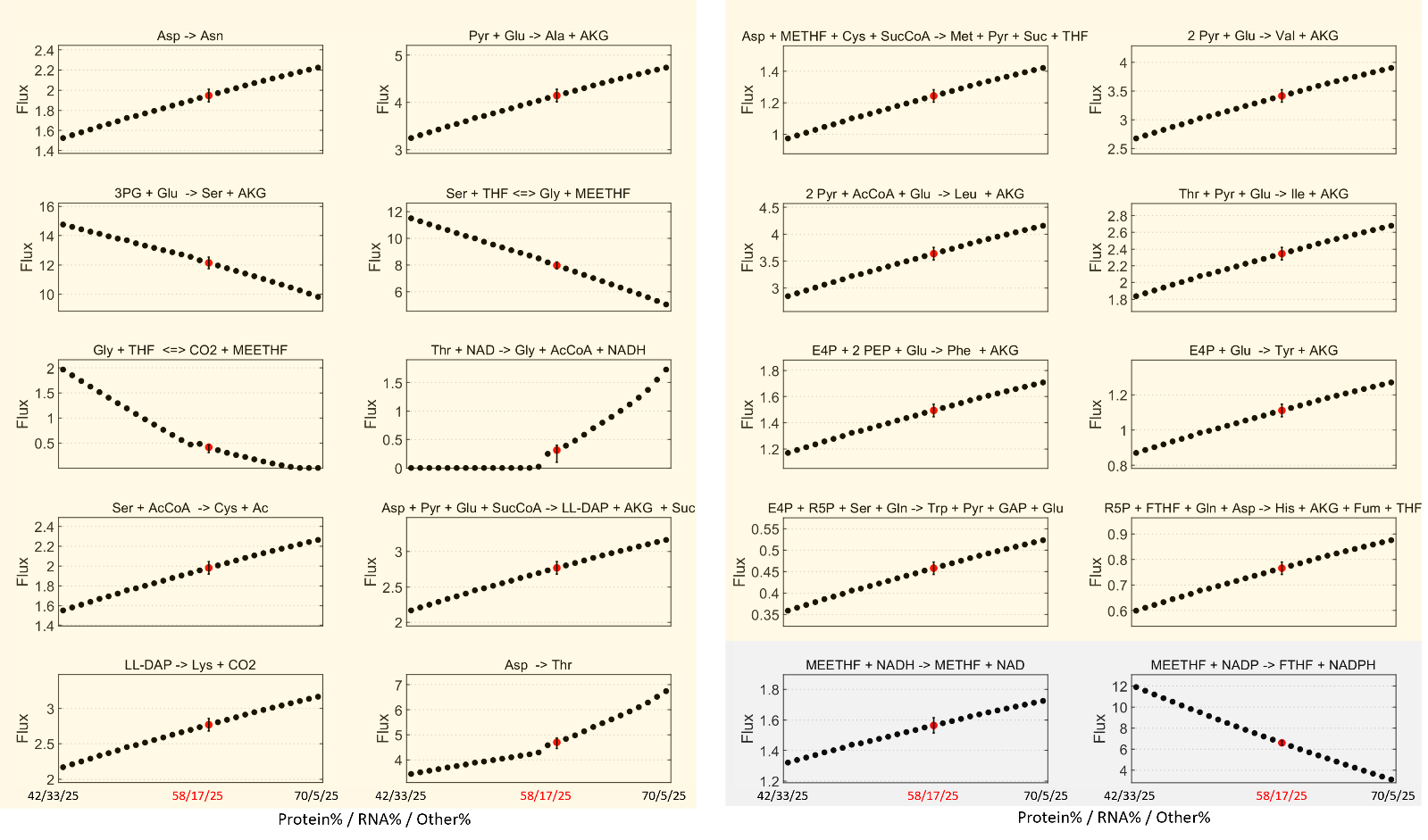

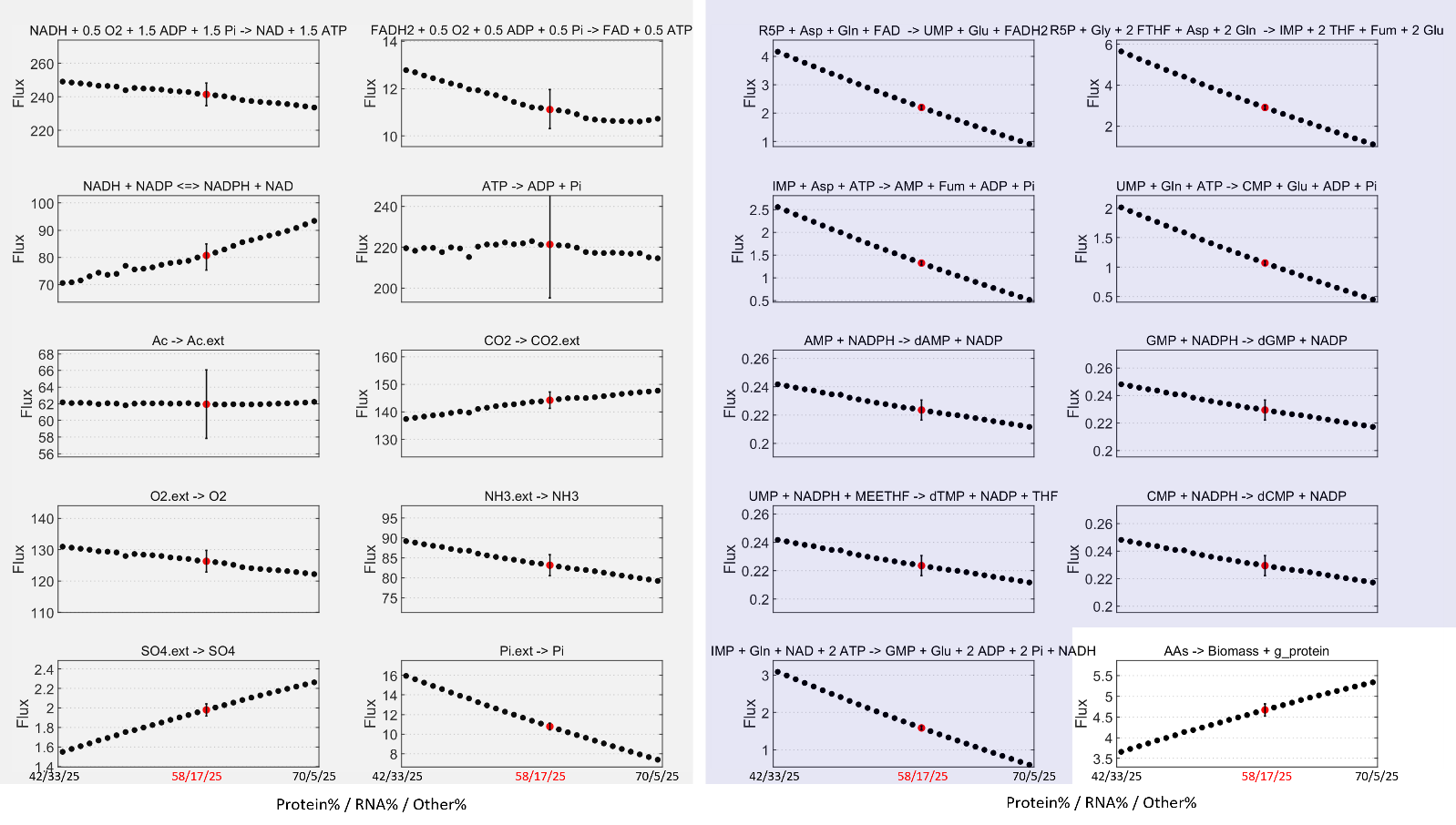

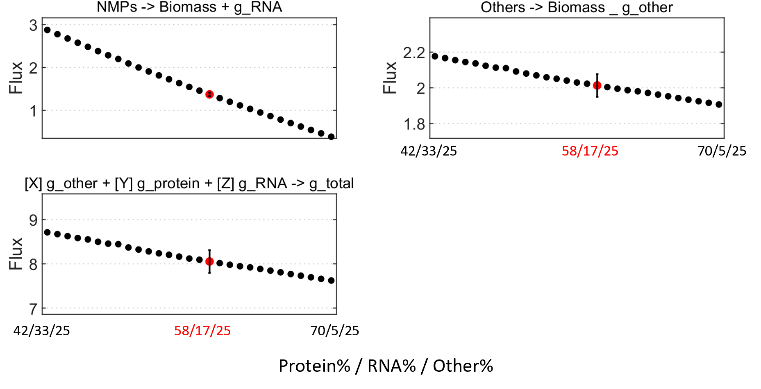

(D) **g_other 30%**

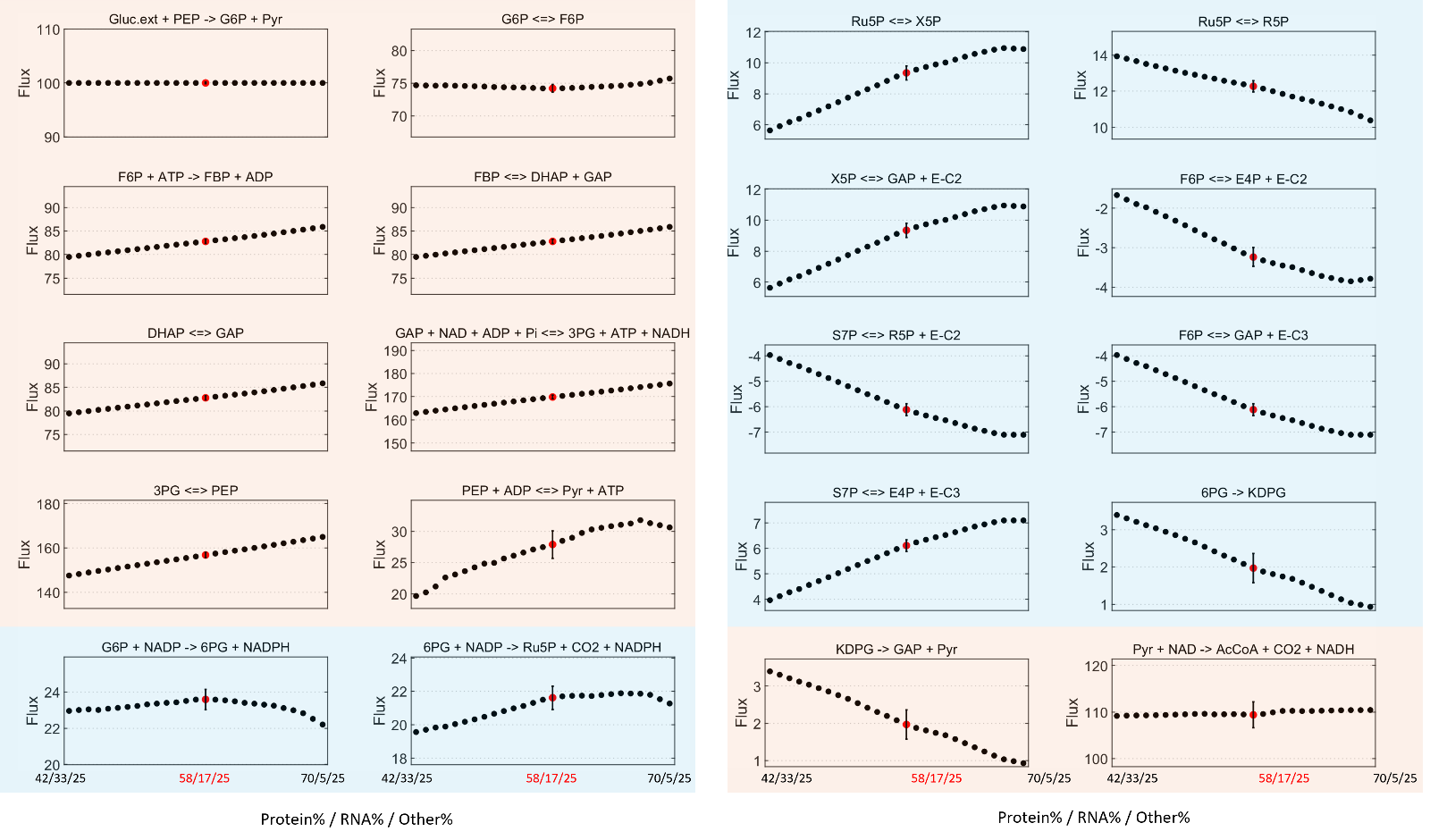

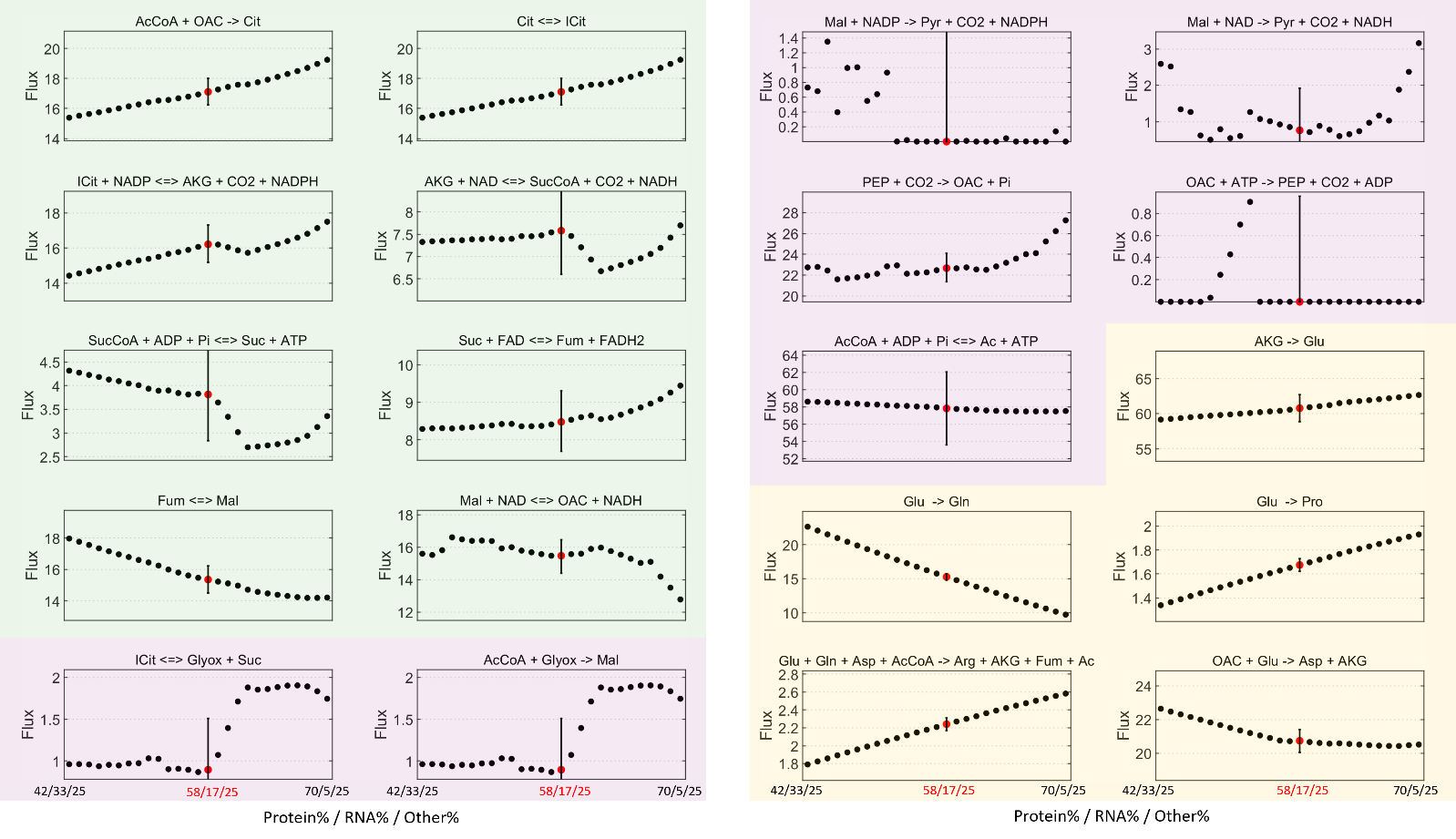

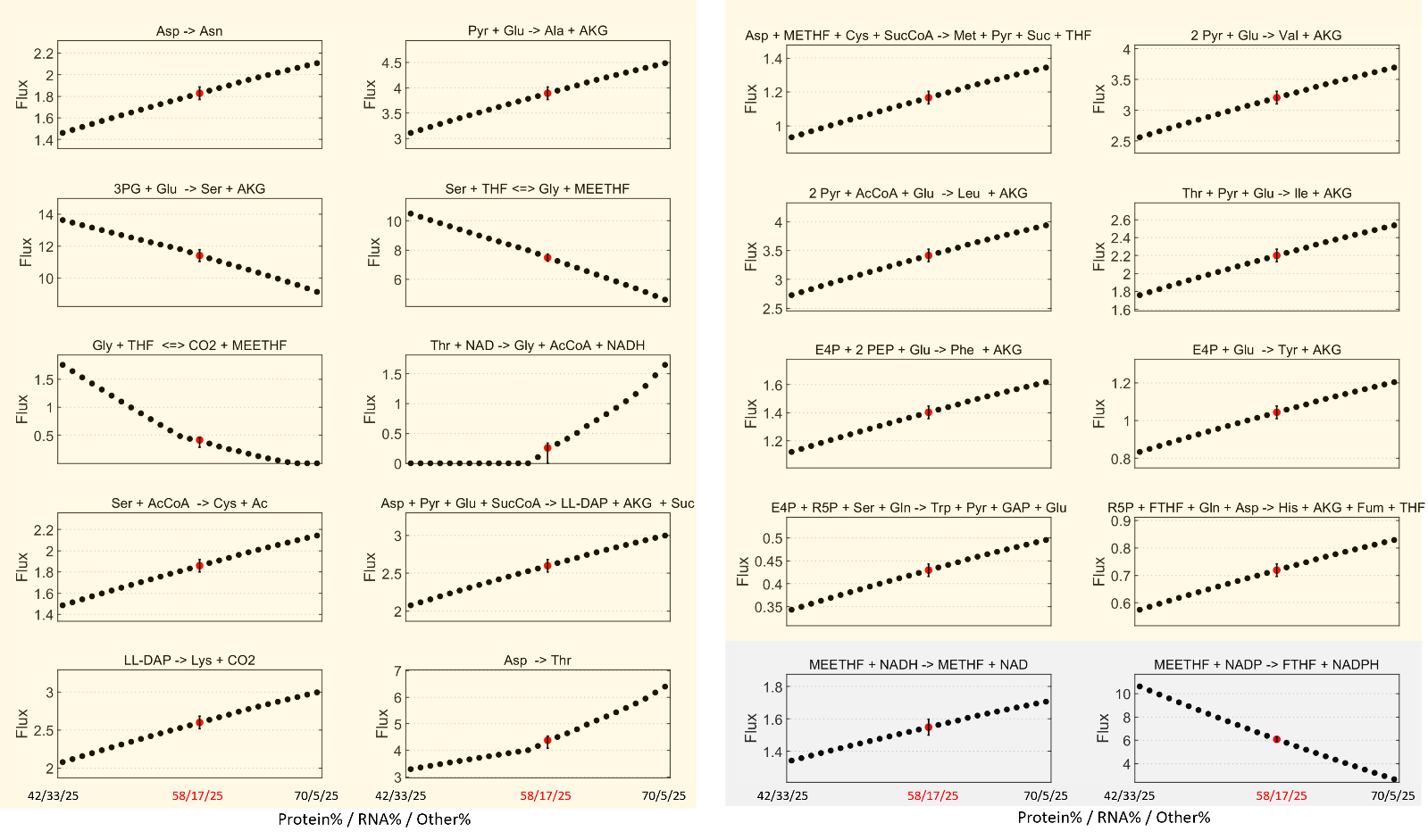

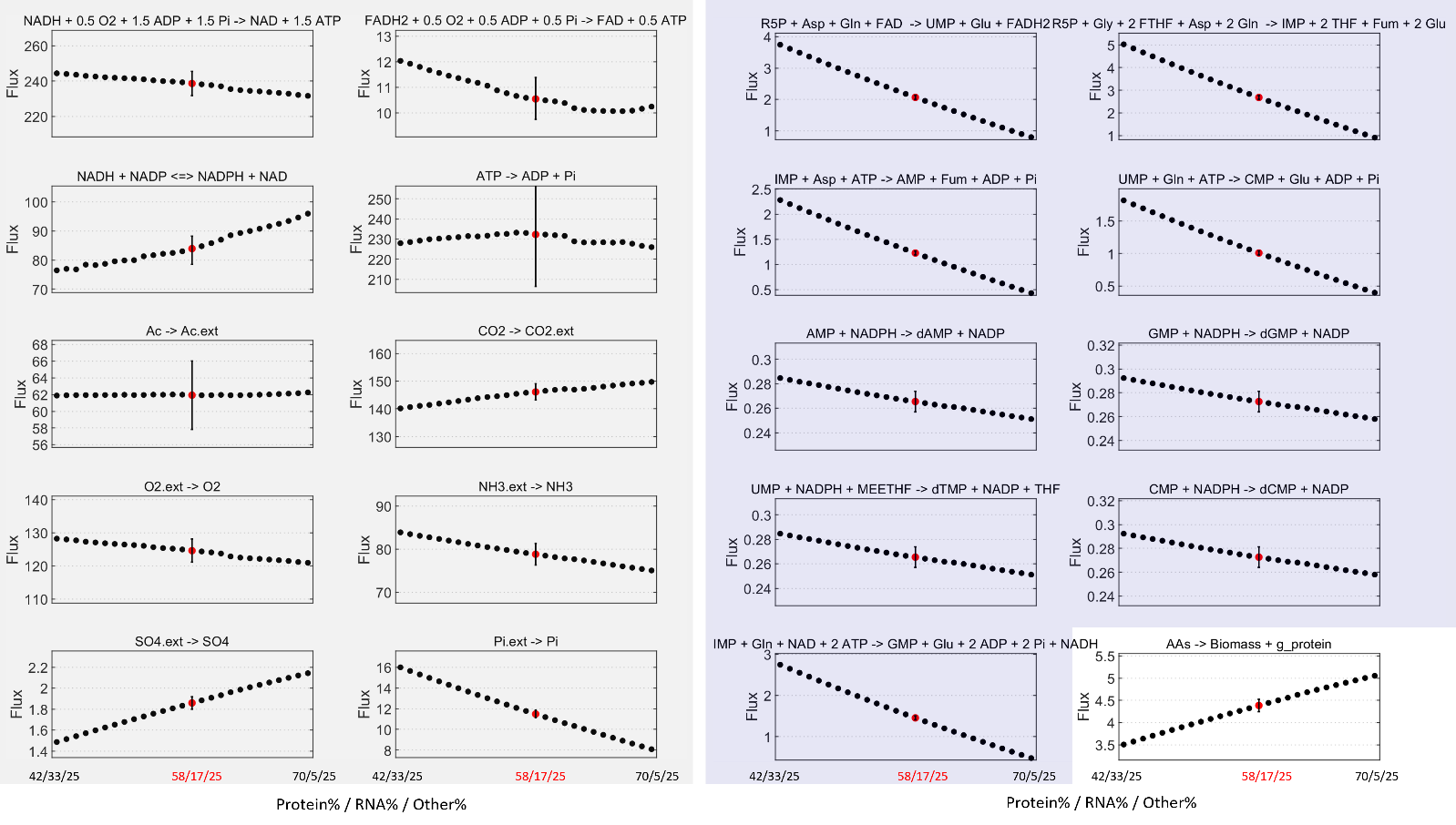

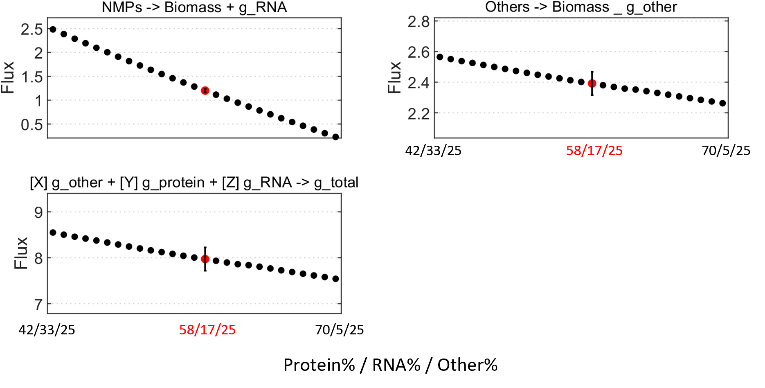

Figure S1: Flux distributions were estimated by ^13^C-MFA and are shown for individual intracellular reactions across different biomass compositions at g_other of 15% (A), 20% (B), 25% (C), and 30% (D). Each subplot represents a single reaction, with black dots indicating flux solutions at different assumed biomass compositions. The red-highlighted point with error bars corresponds to the best-fit solution of each group, with 95% confidence intervals derived from flux statistical analysis. Reactions are color-coded by functional category: glycolysis (red), pentose phosphate pathway (blue), anaplerotic and acetate overflow pathways (purple), tricarboxylic acid (TCA) cycle (green), protein biosynthesis (yellow), nucleotide biosynthesis (dark blue), biomass reactions (white), and other reactions (gray).

**Figure S2: Comparison of flux ranges from Flux Variability Analysis (FVA) and flux estimates from 13C-MFA across biomass compositions**

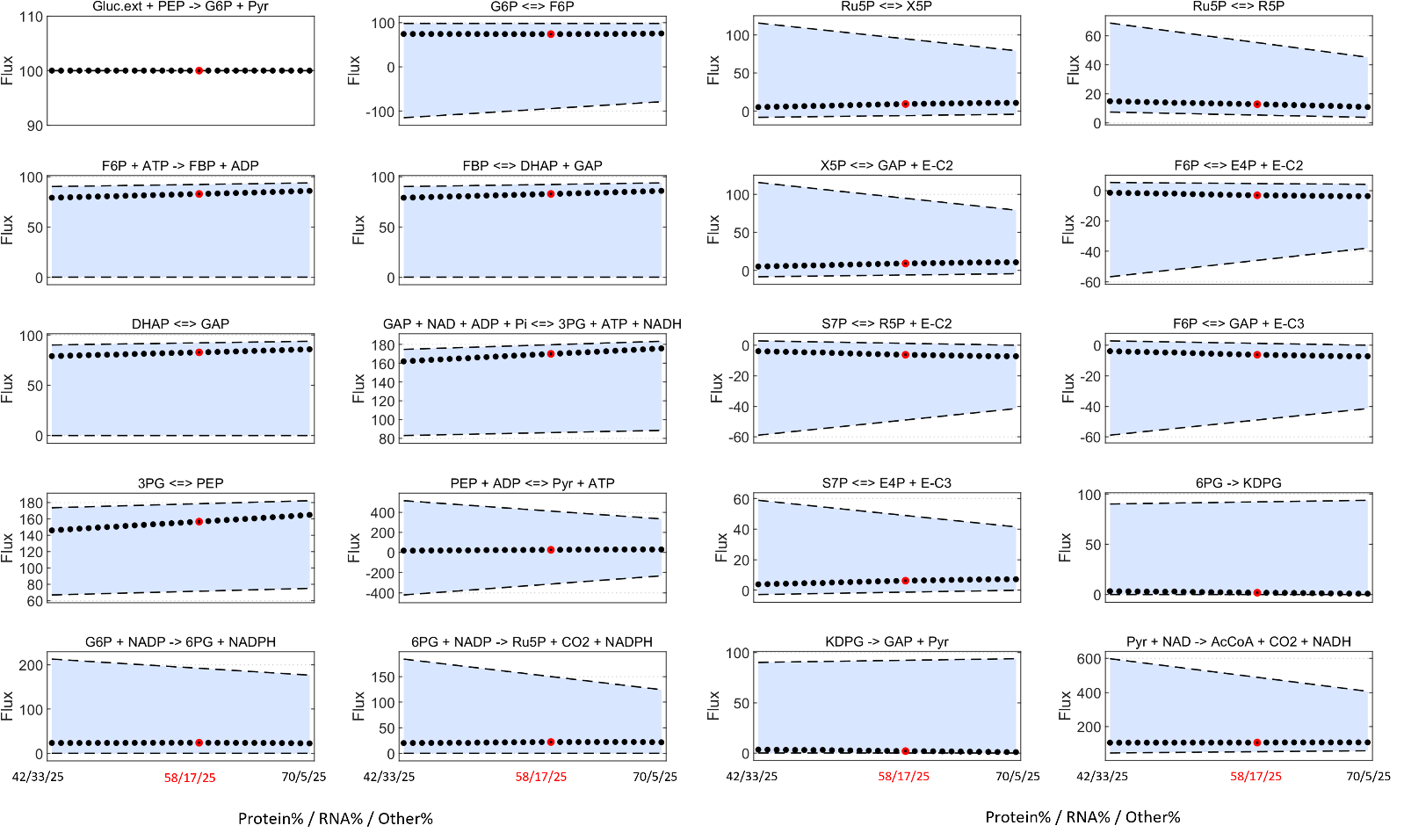

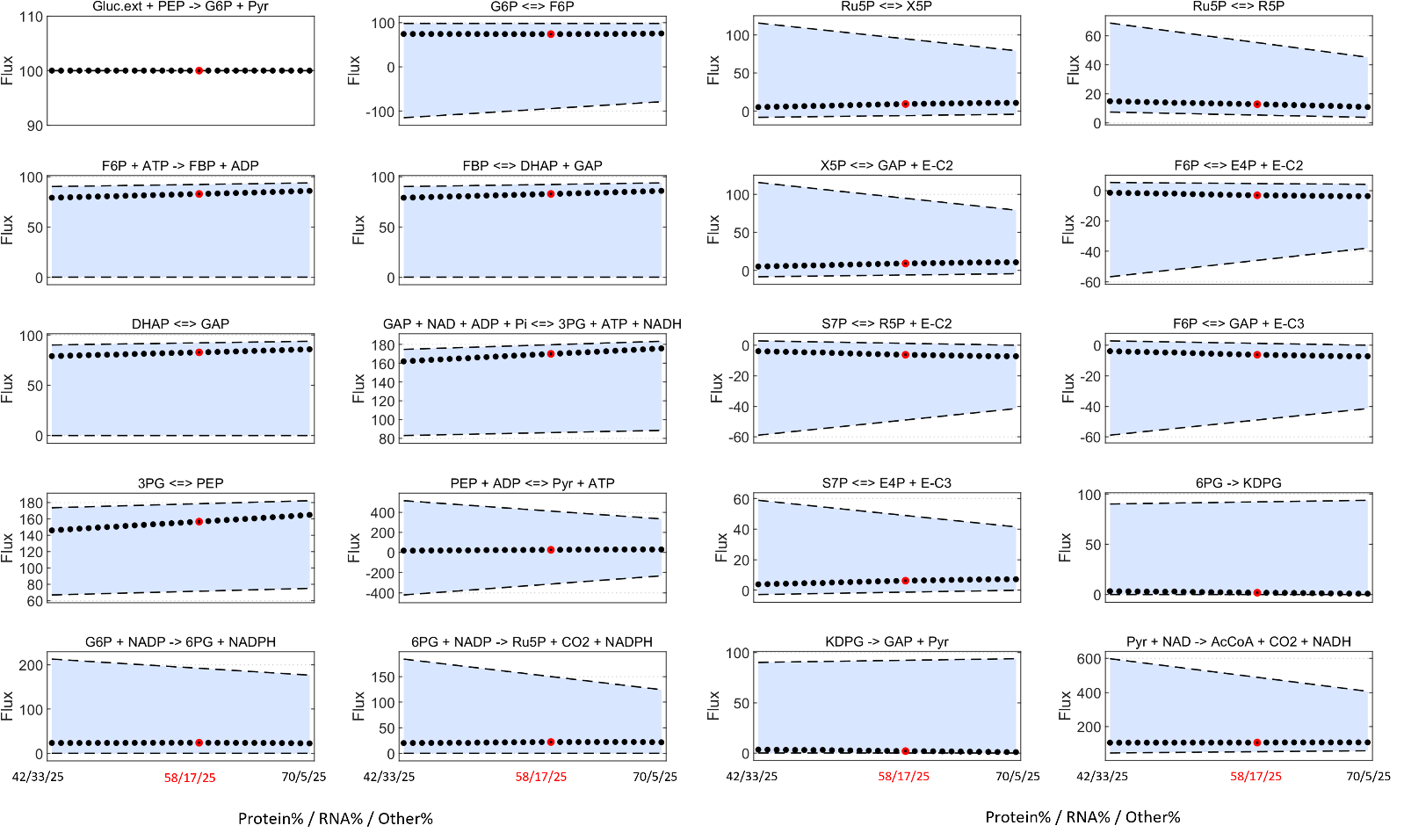

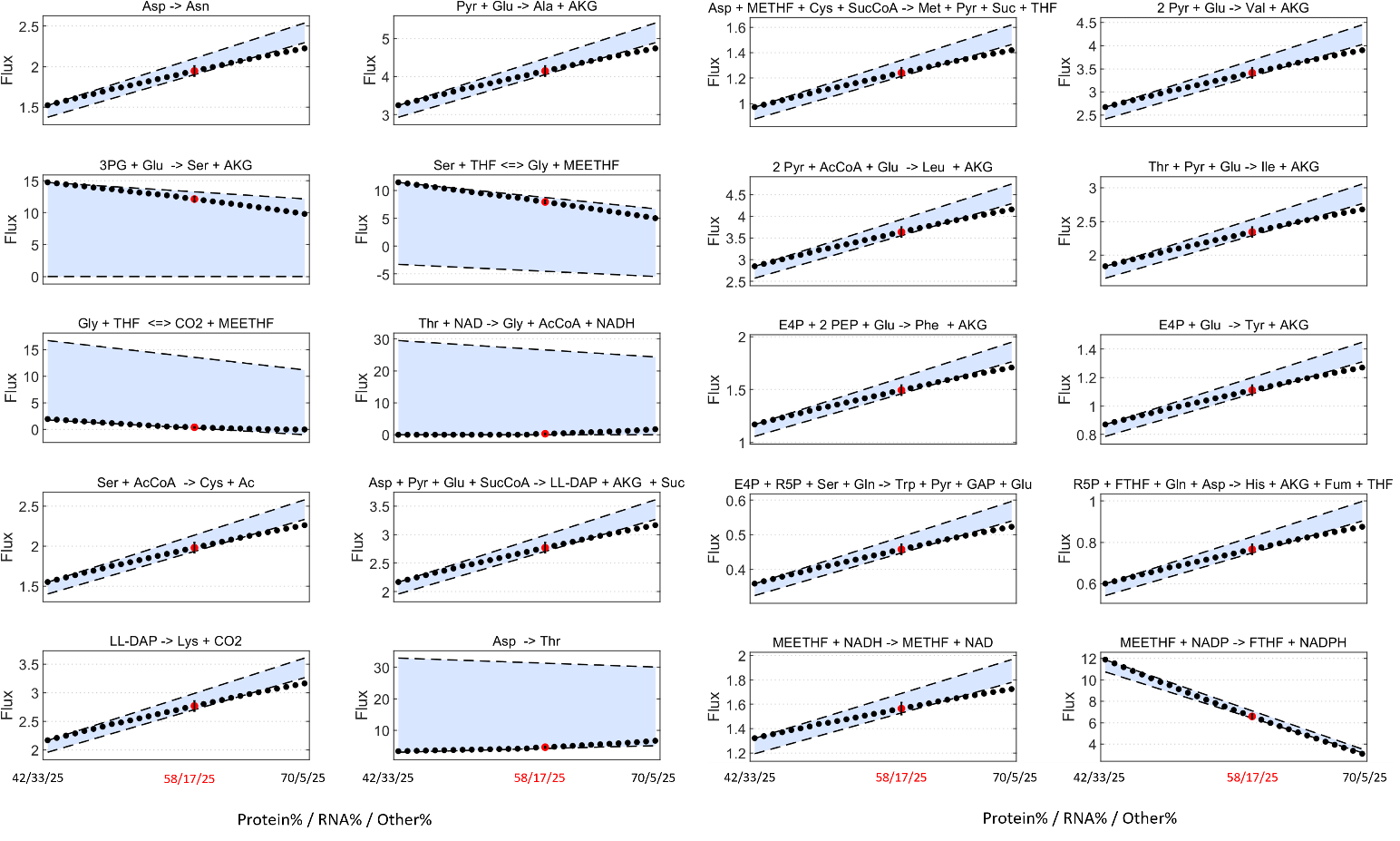

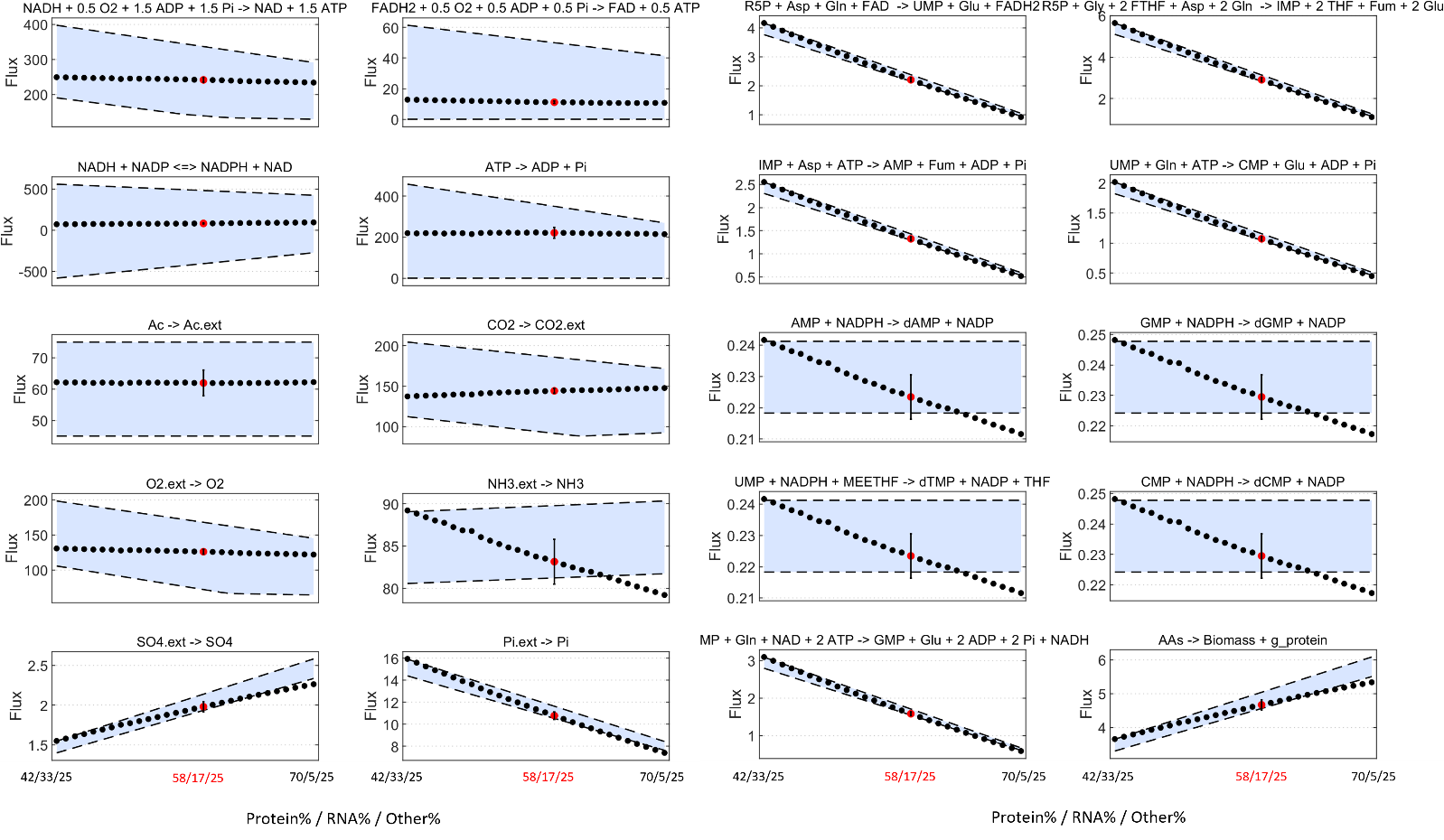

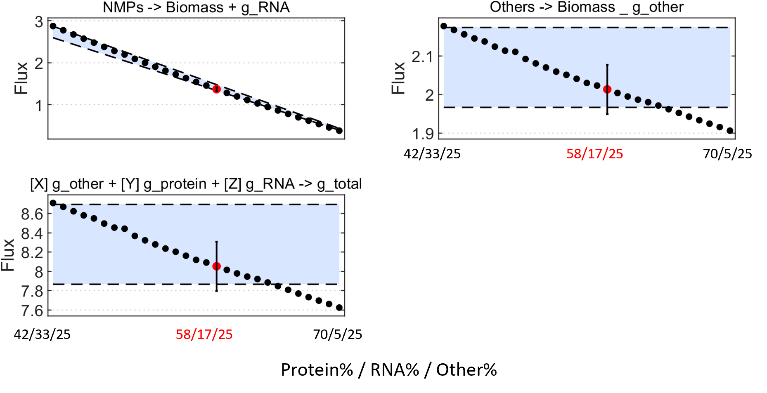

Figure S2:

Each subplot shows a representative reaction from the metabolic network. Black dots represent ^13^C-MFA estimated fluxes under varying biomass compositions (protein and RNA content varied in 1% steps, with g_other fixed at 25%). Red dots indicate the best-fit flux prediction at the composition, yielding the lowest SSR (58% protein / 17% RNA / 25% g_other). Shaded blue regions mark the feasible flux ranges determined by FVA under fixed glucose uptake (100 mmol/gDW/h), constrained acetate production (45–75 mmol per 100 mmol glucose), and biomass yield (0.46 gDW/g ± 5%). Reactions span multiple subsystems, including glycolysis, pentose phosphate pathway (PPP), TCA cycle, amino acid biosynthesis, nucleotide synthesis, and one-carbon metabolism. The FVA range captures the maximum flexibility permitted by stoichiometric and thermodynamic constraints, while ^13^C-MFA further constrains fluxes using isotopic labeling data.
